# Sexual selection purges mutation load, but not overall genetic diversity in populations, decreasing vulnerability to extinction

**DOI:** 10.64898/2026.03.13.711588

**Authors:** Michael D Pointer, Will J Nash, Matthew JG Gage, Tracey Chapman, Alexei A Maklakov, David S Richardson

**Affiliations:** University of East Anglia, Norwich Research Park, Norwich, UK, NR4 7TJ; Earlham Institute, Norwich Research Park, Norwich, UK, NR4 7UZ

## Abstract

Theory suggests sexual selection will enhance population viability by purging deleterious alleles. However, direct genomic evidence for this fundamental idea is scarce and contradictory. We combined long-term experimental evolution with whole-genome re-sequencing to directly test how sexual selection affects mutation load, genomic divergence and extinction risk in *Tribolium castaneum*. After 156 generations, populations evolving under strong sexual selection carried substantially fewer deleterious alleles than populations under weak sexual selection, based on both individual-level estimates of missense and nonsense variants and population-level R_xy_ analyses, indicating more efficient purging of deleterious alleles. In contrast, nucleotide diversity and runs of homozygosity were similar across treatments, indicating that purging was targeted at deleterious variation, and that reduced mutation load in populations under strong sexual selection was not explained by demographic effects. Importantly, population-level mutation load estimates provided best explained extinction risk under inbreeding, directly linking sexual selection to purging and population viability for the first time. Genome scans of high and low sexual selection populations revealed peaks of divergence, enriched for genes involved in courtship, sex discrimination, and seminal fluid proteins. Our results provide direct genomic evidence that sexual selection can reduce mutation load without eroding standing genetic diversity and thus adaptive potential, while driving adaptive divergence in reproductive traits. This beneficial purging may help explain the widespread prevalence of sexual reproduction in nature despite inherent costs and have important ramifications as to how we manage populations of conservation concern.

**Significance Statement:** A long standing and unconfirmed prediction is that sexual selection can improve population health by biasing reproduction away from individuals with high deleterious mutation load. However, direct genomic evidence is lacking, and the indirect data available are scant and contradictory. Now, using whole-genome resequencing of experimental evolved beetle populations, we show that populations evolving under stronger sexual selection carry fewer deleterious mutations and, importantly, are less vulnerable to extinction. Our results provide the first direct genomic support for this long-debated evolutionary process that may help explain the widespread prevalence of sexual reproduction in nature. Our findings also highlight the importance of allowing sexual selection to act in populations of conservation concern.

## Introduction

Sexual selection may be a key mechanism for purging deleterious variation from the genome (Whitlock & Agrawal 2009; MacLellan et al. 2009; Parrett et al. 2022), however, this remains contentious (Arbuthnott & Rundle 2012; Chenoweth et al. 2015; Colpitts et al. 2017). If the mating success of individuals is negatively correlated with the burden of deleterious variants they carry (mutation load; Bertorelle et al. 2022; Grossen and Ramakrishnan 2024), sexual selection will amplify purifying selection via ‘genic capture’ (Rowe and Houle 1996, Tomkins et al. 2004). Theory predicts this process should reduce mutation load in sexual populations relative to asexual ones, and thereby lower extinction risk, particularly under inbreeding when recessive load is expressed (Agrawal 2001; Siller 2001; Whitlock and Agrawal 2009). Several studies have used experimental approaches to address this question with mixed results (Radwan 2004, Jarzebowska and Radwan 2009, McGuigan et al. 2011, Almbro and Simmons 2014, Lumley et al. 2015, Prokop et al. 2019, Grieshop et al. 2021, Parrett et al 2022, Leigh et al. 2025), but none directly test the effect of sexual selection on functionally characterised variation. Recent work suggests that sexual selection does not affect overall mutation load, thus questioning the role of sexual selection in purging of deleterious alleles (Leigh et al. 2025). Here we provide the first test of this theory that directly quantifies functionally annotated deleterious and loss-of-function variants in populations evolving with or without sexual selection and explicitly links this genomic load to extinction risk.

Experimental evolution studies have reported indirect evidence that sexual selection appears to slow the accumulation of deleterious mutations or improves non-sexual fitness components (Radwan 2004; McGuigan et al. 2011; Almbro and Simmons 2014; Lumley et al. 2015; Cally et al. 2019; Grieshop et al. 2021; Parrett et al. 2022), or, alternatively, that it yields little net fitness benefit and can exacerbate sexual antagonism (Hollis and Houle 2011; Arbuthnott and Rundle 2012; Chenoweth et al. 2015; Prokop et al. 2019; Leigh et al. 2025). However, these studies infer mutation load indirectly from changes in phenotypes or genome-wide diversity, rather than from direct estimates of functionally annotated deleterious variants. Indirect evidence consistent with the purging of load under sexual selection comes from phenotypic studies in *Tribolium castaneum* flour beetles. These studies inferred a reduction in load because populations evolved under elevated sexual selection showed evidence of increased performance across fitness-related traits (Lumley et al. 2015; Dugand 2018; Godwin et al. 2020). For example, beetle lines under strong (versus weak) sexual selection showed increased resistance to extinction under inbreeding (Lumley et al. 2015). No study has yet directly quantified mutation load in experimental lines evolved under differing levels of sexual selection which is the major gap we address here.

We functionally annotated genomic variants to directly quantify the burden of deleterious sites as an estimate of mutation load in 84 individual genomes. These genomes were sequenced from the six *T. castaneum* lines described in Lumley et al. (2015) that had been experimentally evolved for 156 generations under either monandrous (absent / weak sexual selection) or polyandrous (strong sexual selection) mating regimes. We directly determined genome-wide nucleotide diversity and estimated mutation load by categorising mutations into functional categories (missense tolerated, missense deleterious, nonsense). We also determined relative mutation load at the population-level by comparison of allele frequencies across these different functional categories of variants. Alongside this, we identified loci where allele frequency differences were significantly associated with sexual selection regime, providing putative targets of selection across the genome. Using these loci to define highly differentiated regions of the genome, we compared signatures consistent with the functional impacts of purifying selection, identified in our earlier analyses, and those associated with the action of directional selection. In addition, using data on survival under inbreeding from the same lines (Lumley et al. 2015) we tested for associations between mutation load and extinction probability. Importantly, this allowed us to test whether purging of load in high sexual selection lines is associated with their observed inbreeding resilience (Lumley et al. 2015; Godwin et al. 2020).

Our results showed a striking elevation in the purging of genome-wide deleterious variation in response to sexual selection. This was not associated with any reduction in overall genetic diversity and so cannot be explained by demographic differences, such as differential inbreeding among the sexual selection regimes during experimental evolution. Instead, elevated sexual selection has resulted in the specific elimination of deleterious genetic variation itself. Importantly, it is the overall genome-wide load of such deleterious variants that remain after the selection regime that then best predicts population extinction risk in this experiment. In addition, genomic windows which diverge between replicated lines were explained by positive selection for beneficial alleles with predicted roles in reproductive biology, courtship behaviour, sex discrimination, and male seminal fluid protein function.

## Results

### Sexual selection is associated with a lower burden of deleterious variants

We estimated mutation load in lines of *Tribolium castaneum* beetles drawn from populations evolved under experimental regimes of strong or weak sexual selection for 156 generations (Lumley et al. 2015). Briefly, these regimes either allowed polyandry, 5 males to 1 female, elevating the potential for SS; or enforced monandry, 1 male to 1 female, minimising / removing the potential for SS (while controlling for population size (see methods). Individual mutation load was estimated by quantifying variants in three functional categories: missense tolerated, missense deleterious, and nonsense. Variant counts were normalised by the number of synonymous variants in the same individual to obtain a ratio, accounting for variation in genetic diversity and sequencing coverage. These normalised per-individual counts of high-quality SNPs provide a relative measure of functional mutational burden, which we use as an estimate of mutation load in population-level comparisons.

Considering the coding regions of autosomes, the ratio of missense tolerated variants relative to synonymous variants, per individual, was not significantly different between monandrous and polyandrous populations (β = −1.65 x 10^-5^, SE = 6.06 x 10^-4^, p = 0.98; figure 1A). For missense deleterious variants, ratios were significantly lower in polyandrous populations than in monandrous ones (β = −1.08 x 10^-3^, SE = 1.15 x 10^-4^, p < 0.0001). Populations evolved under polyandry carried fewer nonsense variants than those evolved under monandry (β = −5.23 x 10^-4^, SE = 3.98 x 10^-5^, p < 0.0001).

**Figure 1.**
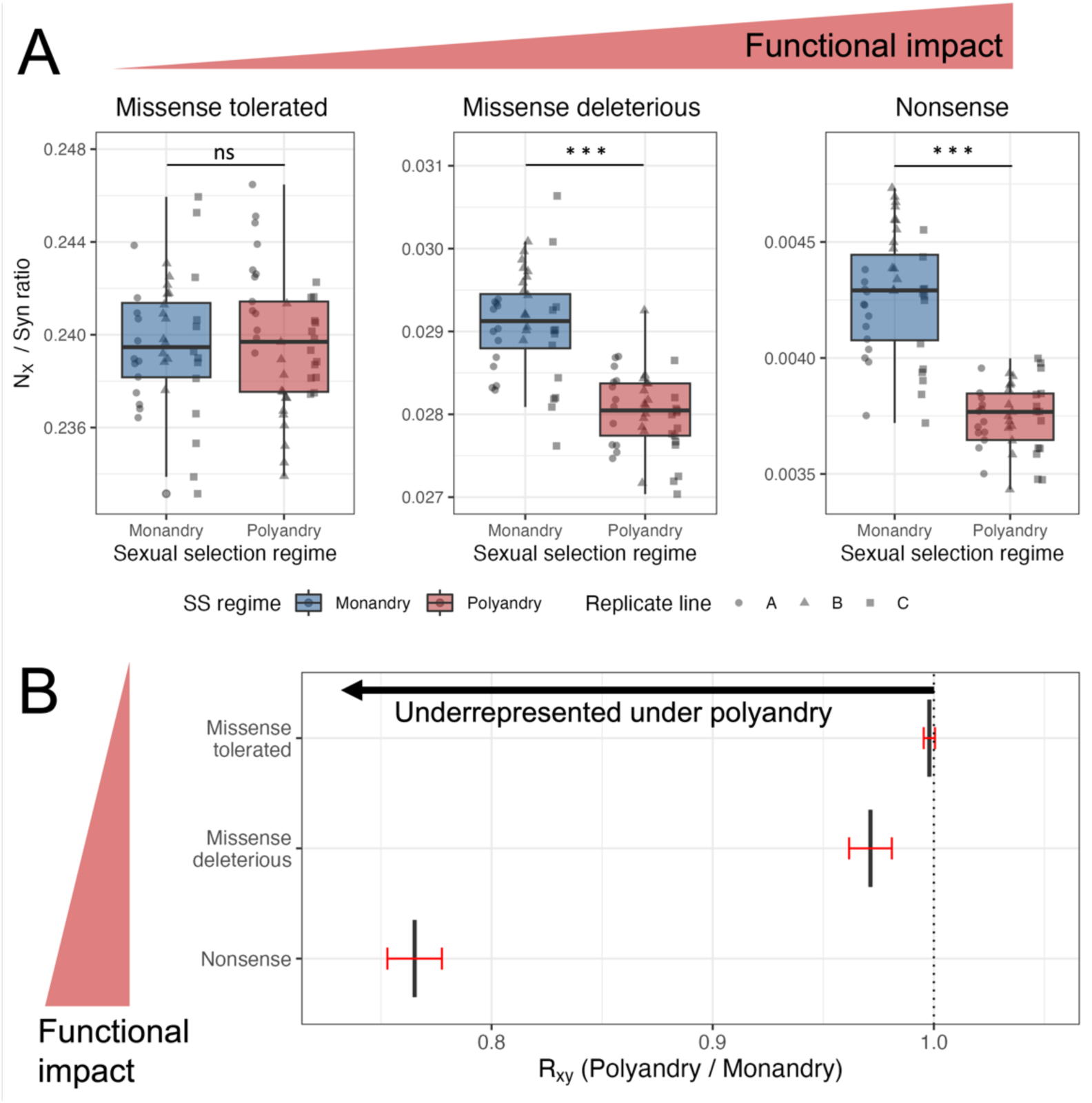
Strong sexual selection is associated with a significantly lower burden of deleterious variants. Genetic variation categorised by increasing deleterious functional impact (red wedges) in populations of *Tribolium castaneum* evolved under either monandrous or polyandrous sexual selection regimes for 156 generations. A) The ratio of missense tolerated, missense deleterious and nonsense (N_X_) sites to synonymous sites, per individual. B) R_xy_ analysis contrasting allele frequencies in monandrous and polyandrous populations, across functional categories of high-quality SNPs, normalized by R_xy_ of intergenic variants. R_xy_ < 1 indicates a relative frequency deficit of the corresponding category under polyandry. Red bars represent 95% CIs computed from block jack-knife standard errors.

Across autosomal loci, we found no evidence that the effect of sexual selection regime differed between males and females for any of the three functional variant categories (interaction term: p > 0.53). After removing the interaction term, sex did not predict the relative number of autosomal missense tolerated, or missense deleterious variants (p > 0.18; figure S2). In contrast, sex was associated with the number nonsense variants, with females carrying fewer than males (β = −1.38e^-4^, SE = 3.63 x 10^-5^, p < 0.01). *T. castaneum* has an X/Y sex determination system with males the heterogametic sex (Juan and Petitpierre 1991). On the X chromosome, the ratio of missense tolerated or missense deleterious variants to synonymous variants did not vary between sexual selection treatments (p > 0.07; figure S3). However, polyandrous populations exhibited marginally more nonsense variants than monandrous populations (β = 2.37×10^-4^, SE = 1.16 x 10^-4^, p = 0.044). As on the autosomes, there was no evidence for differing effect of sexual selection regime on males and females across the three variant categories tested using the X chromosome (all p > 0.21). With the interaction term removed, sex was significantly associated with the relative number of missense tolerated variants, with lower counts in males (β = −1.09 x 10^-2^, SE = 2.03 x 10^-3^, p < 0.0001; figure S4). A marginally significant association with sex, following the same pattern, was also observed for missense deleterious variants (β = −8.09 x 10^-4^, SE = 4.01 x 10^-4^, p < 0.04) but there was no difference in the relative number of nonsense variants (p = 0.94).

To evaluate mutation load at population level, we compared the frequency of missense tolerated, missense deleterious, and nonsense variants between sexual selection regimes. For this analysis line replicates within each regime were pooled to give two ‘populations’ to contrast, to obtain stable estimates of allele frequencies for each functional class and to characterise regime-level differences in the genomic distribution of functional variants. We computed R_xy_ (described by Xue et al. 2015; after the method of Dussex et al. 2023) for each functional category between monandrous and polyandrous populations, where values <1 indicate that variants of the focal category are underrepresented in polyandrous populations relative to monandrous populations. Missense tolerated R_xy_ confidence intervals (CIs) crossed one, indicating no mean difference in frequency between sexual selection regimes (figure 1B). Missense deleterious variants both showed R_xy_ values close to one but with CIs not including one, suggesting a small deficit of these variants under polyandry. Nonsense variants had the lowest values of R_xy_ (jackknife mean = 0.77) with CIs excluding one, indicating that variants with highly deleterious functional impact are strongly underrepresented in genomes from the polyandrous regime (figure 1B). The pattern of R_xy_ across functional categories observed on the autosomes was not replicated on the X chromosome (figure S5). We found no evidence for a relationship between functional impact and purging strength on the X chromosome, though our ability to detect an effect was likely impacted by small numbers of variants on the X, especially in high impact categories.

### Survival of sexual selection lines following inbreeding is more parsimoniously explained by mutation load than by sexual selection regime

Higher mutation load is predicted to increase the risk of population extinction under inbreeding. Lumley et al. (2015) showed that polyandrous populations had lower extinction vulnerability than monandrous populations, under enforced inbreeding, using the same experimental lines we employ in this study. To explore the link between mutation load and extinction vulnerability, we extended the survival analysis framework of Lumley et al. (2015) to include our line-level genomic estimate of mutation load. Replicating their findings, sexual selection regime significantly predicted time to extinction, with polyandrous populations exhibiting lower extinction risk (table 1). However, AIC model comparison showed strongest support for a model including mutation load alone (AIC = 276; AIC weight = 0.66; table 1), compared to a model including sexual selection regime only (AIC = 280). When both predictors were included, mutation load remained marginally significant (p = 0.046), whereas the effect of sexual selection regime was non-significant (p = 0.90; AIC = 278). This drop in significance is expected, given that we know sexual selection regime and mutation load to covary, but is further evidence that mutation load provides explanatory power beyond that captured by sexual selection regime.

**Table 1.** Survival under inbreeding is better explained by mutation load than by sexual selection regime. Functional characterisation of genomic variants was used to estimate mutation load in 84 genomes sequenced from *Tribolium castaneum* lines evolved under strong (polyandry) or weak (monandry) sexual selection (SS) regimes for 156 generations. Three mixed models (GLMM) were fitted to population survival data, when families derived from these same lines where subjected to enforced inbreeding (from Lumley et al. 2015), modelling survival as a product of mutation load or SS regime or both. Mutation load alone better explained survival than did SS regime, or a combined model.

| Model | SS regime effect<br>(coef, p) | Mutation load effect<br>(coef, p) | LokLik | AIC | AIC weight |
| --- | --- | --- | --- | --- | --- |
| SS regime | -0.730, p = 0.026 | – | -138.0 | 280.0 | 0.087 |
| Mutation load | – | 299.5, p = 0.003 | -136.0 | 276.0 | 0.667 |
| SS regime + mutation load | 0.065, p = 0.900 | 314.8, p = 0.046 | -136.0 | 278.0 | 0.245 |

### Sexual selection does not affect overall genome-wide variation

We found no evidence that genome wide nucleotide diversity (π) differed between monandrous (0.153 ± 0.026) and polyandrous (0.168 ± 0.023) mating regimes (linear model, F₁,₄ = 0.19, p = 0.68). The genomic inbreeding coefficient (F_ROH_), computed, per individual, as the proportion of the callable genome in runs of homozygosity (>0.2Mb), did not differ between monandrous and polyandrous regimes (0.024 ± 0.002 vs. 0.026 ± 0.001 respectively, β = 1.48 x 10^-3^, SE = 3.13 x 10^-3^, p = 0.662; figure 2A). The distribution of runs of homozygosity of different lengths also appeared similar between sexual selection regimes (figure 2B).

**Figure 2.**
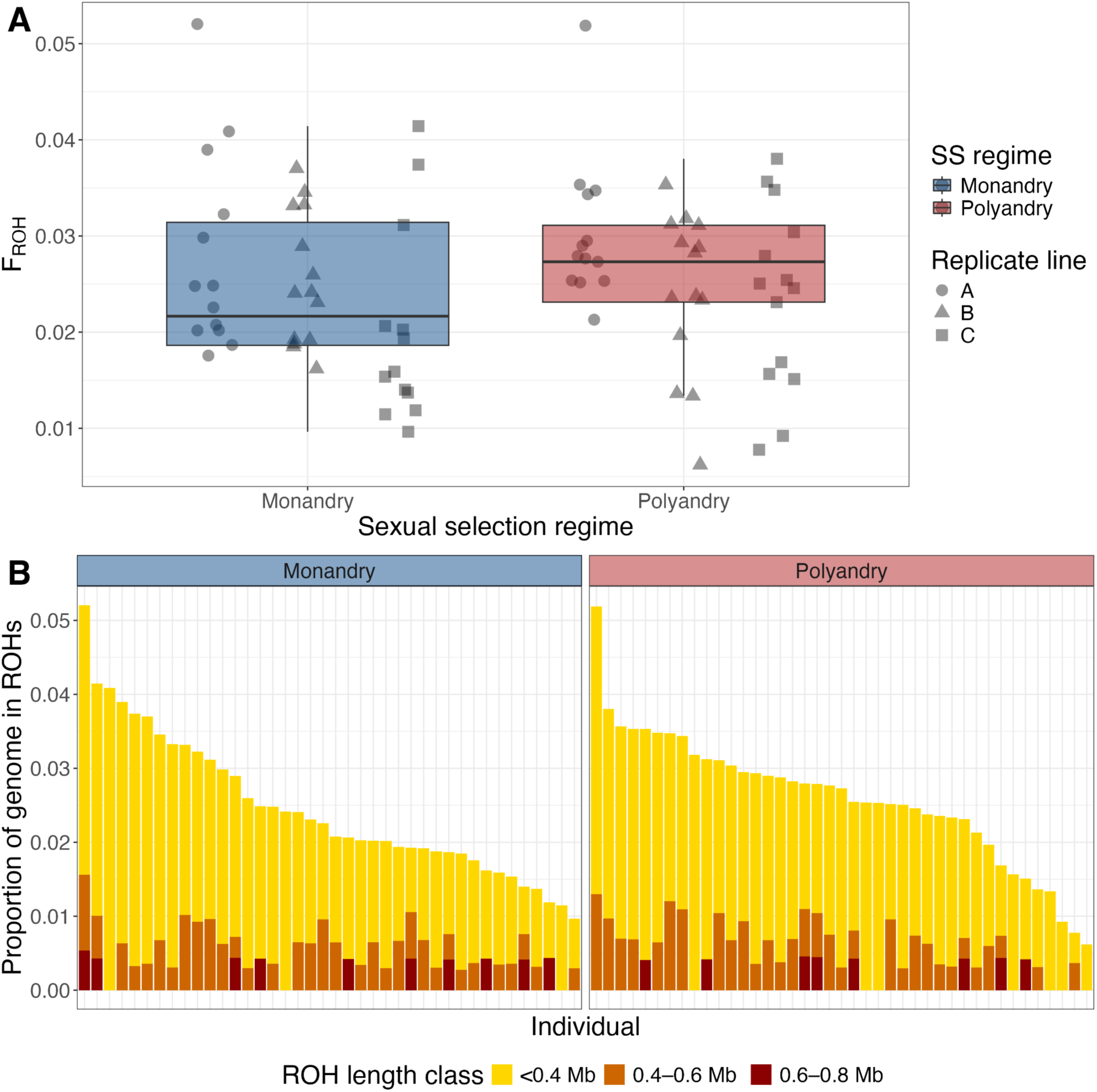
Sexual selection does not affect individual inbreeding. Inbreeding assessed via runs of homozygosity (ROH) in the genomes of *T. castaneum* experimentally evolved under monandrous and polyandrous sexual selection (SS) regimes. **A**) genomic inbreeding coefficients, proportion of the genome present in ROH > 0.2Mb. No significant difference between monandrous and polyandrous populations (LMM: β = 1.48 x 10^-3^, SE = 3.13 x 10^-3^, p = 0.662). **B**) The proportion of the genome present in ROH of different lengths in each individual.

### Peaks of genetic divergence between populations evolved under high or low sexual selection

To identify genomic regions that had significantly diverged following 156 generations of experimental evolution under the two sexual selection regimes we performed a genome-wide scan for SNP-wise allele frequency differentiation, using BayPass (Olazcuaga et al. 2020). We identified 216 autosomal and 226 X chromosome SNPs as C2 outliers, grouped into 15 divergent windows genome-wide (figure 3; table S4). Outlier SNPs on the X chromosome clustered into two strong peaks, separated by ∼200kb (figure 3B). Returning the closest gene to each of the 442 total candidate SNPs gave a set of 54 genes (table S3). Considering only genes located within 10kb of just the top C2 SNP from each of the 15 candidate regions, 39 genes were identified (table 2), with 26 of these having well characterised homologs in *Drosophila melanogaster*.

**Figure 3.**
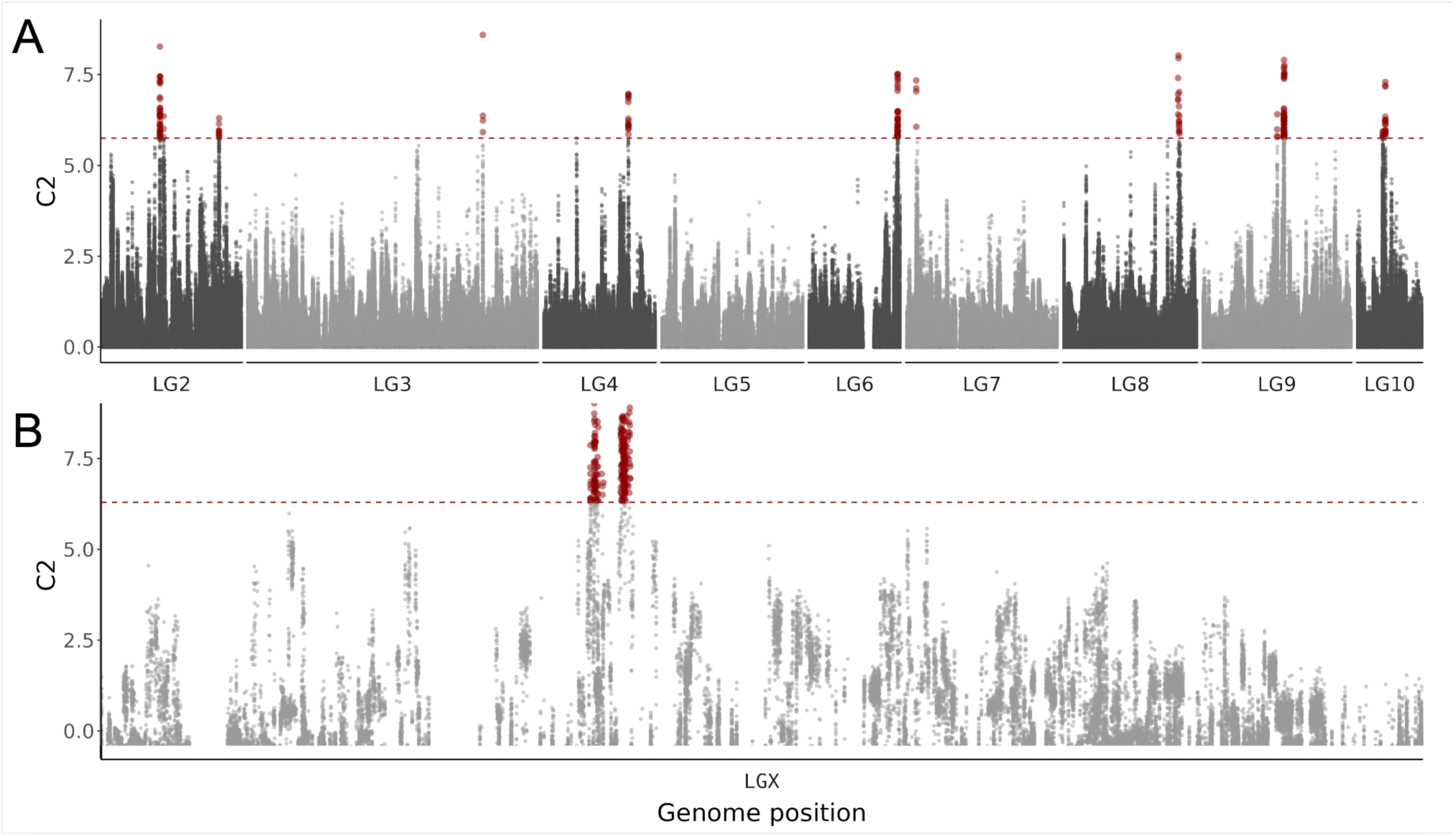
Genomic loci showing allele frequency differentiation associated with sexual selection regime in experimentally evolving *Tribolium castaneum* populations. Values of the C2 statistic generated by BayPass analysis (Olazcuaga et al. 2020) measuring SNP-wise associations with the two sexual selection regimes (polyandrous vs. monandrous) while accounting for shared population history. Red points are in excess of the 0.999 quantile of C2 values from a BayPass run on a pseudo-observed dataset of putatively neutral SNPs, shown by the dotted line. Separate analyses were performed for **A)** autosomal linkage groups, and **B)** the X chromosome (LGX).

**Table 2.**
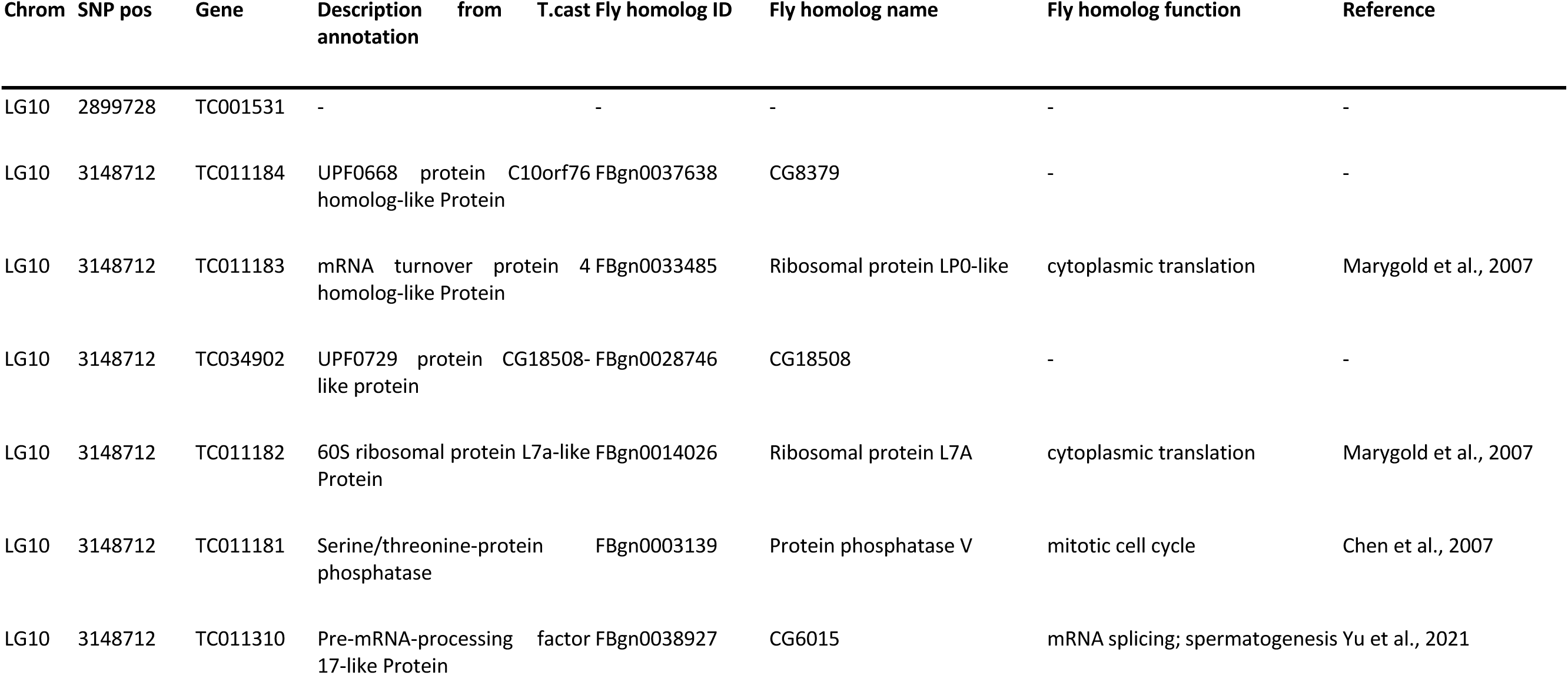
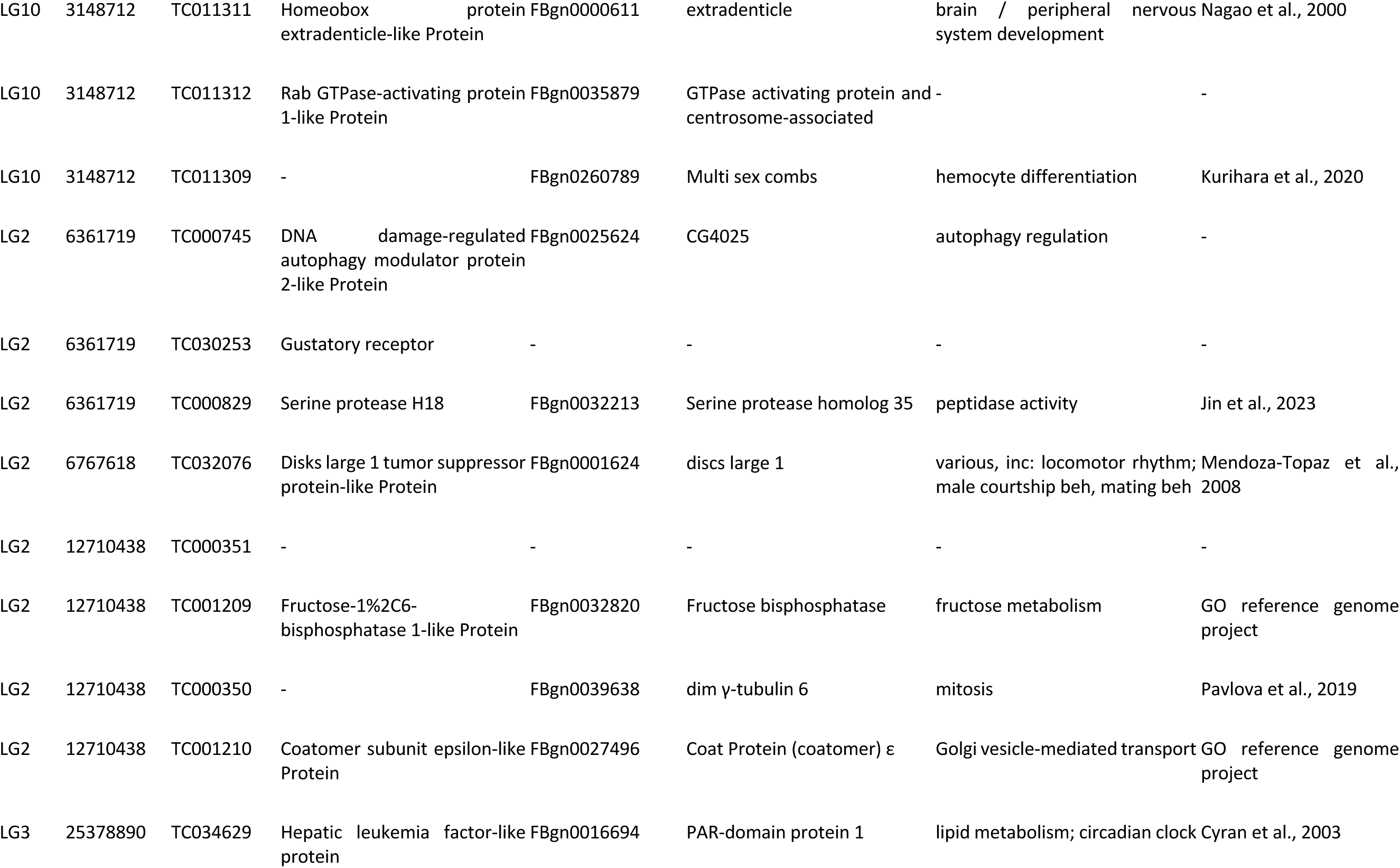
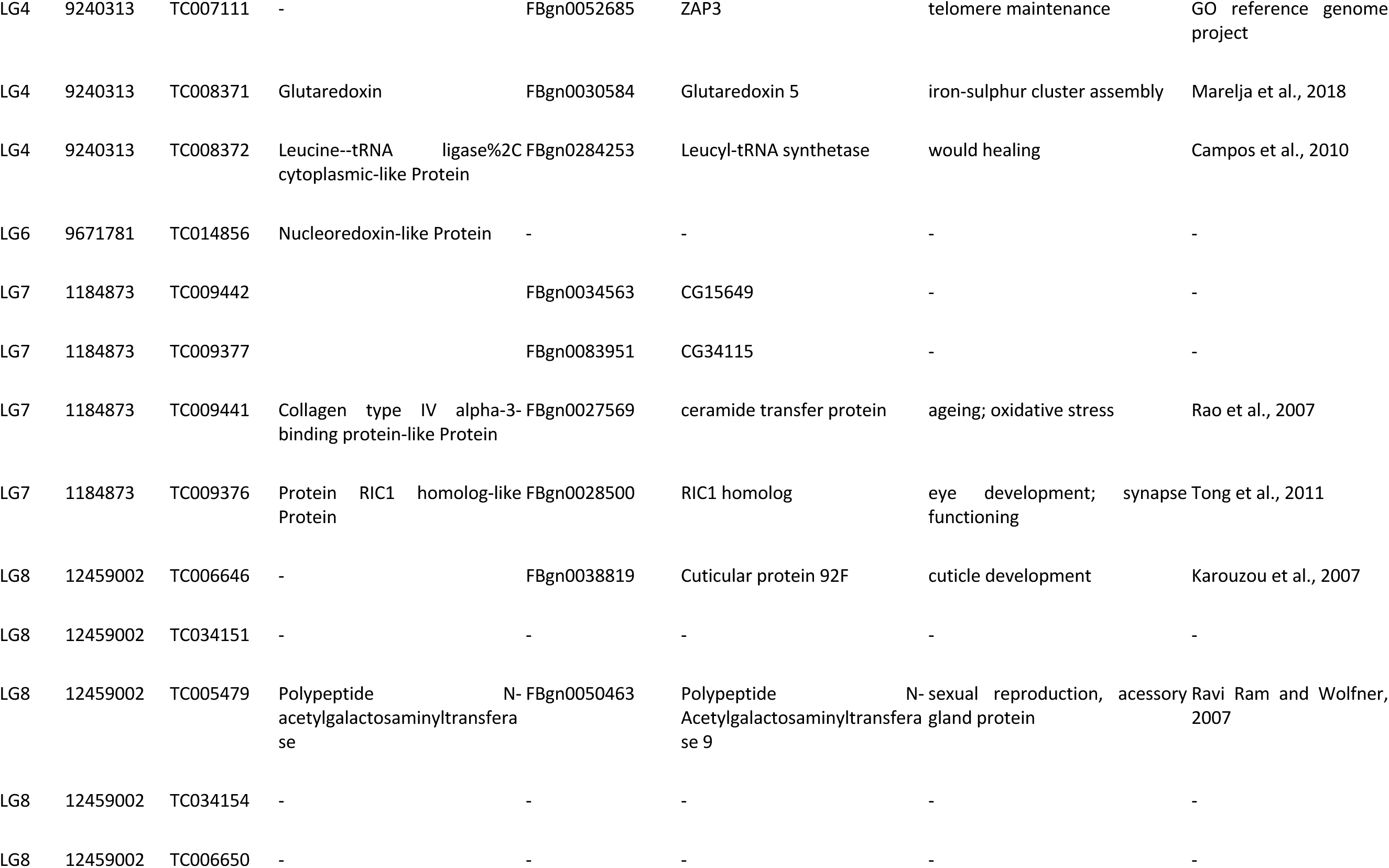
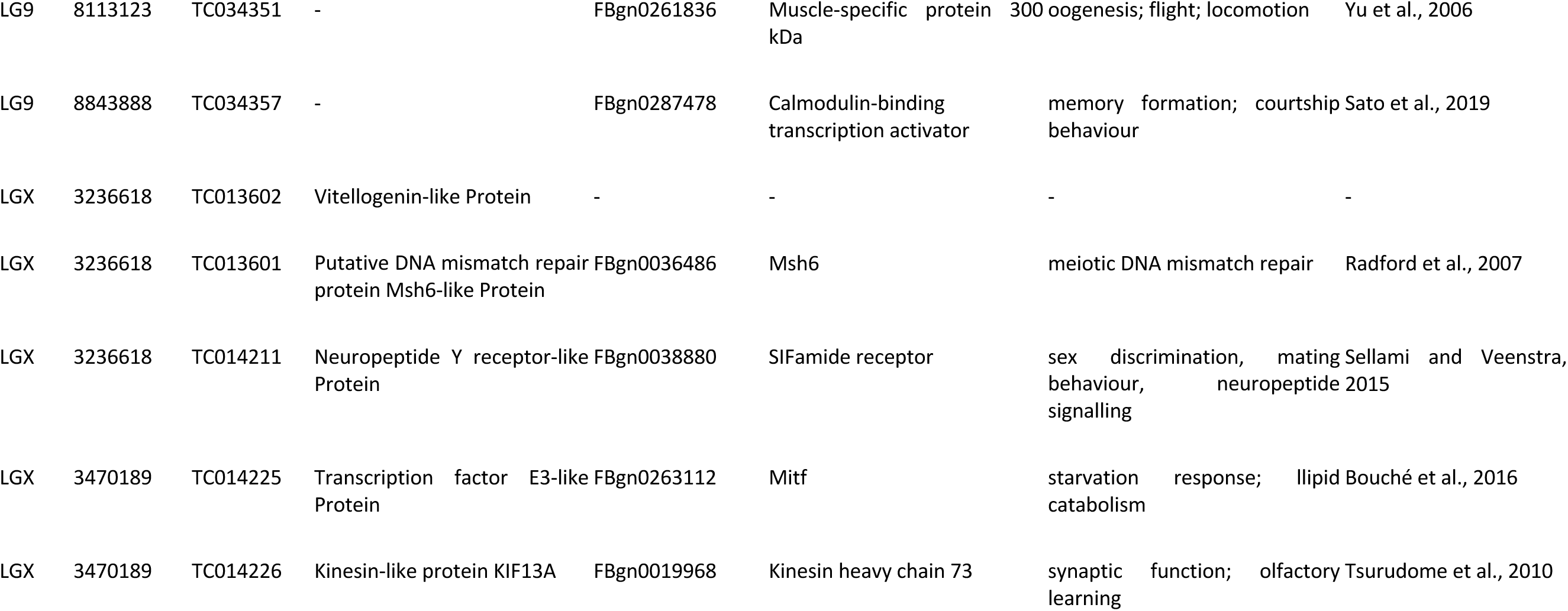
Genes from significantly divergent regions. Genes identified in regions divergent between *Tribolium castaneum* individuals evolved under either strong or weak sexual selection. The genome scan for divergence was performed with BayPass () between three polyandrous and three monandrous populations evolved under these regimes for 156 generations. Gene functions were obtained from Tcas5.2 genome annotation and cross-referenced with FlyBase via the iBeetlebase database.

Although neither gene set returned was enriched for gene ontology terms (G:profiler; Kolberg et al. 2023), background: all Tcas5.2 genes; p > 0.05), the 26 genes with functional information (from either the Tcas5.2 annotation or *D. melanogaster* homologues) reflected a diverse range of functions (table 2) including protein synthesis, lipid metabolism and oxidative stress tolerance. Ten genes also had known functions (from the Tribolium 5.2 annotation or from Drosophila homologues) directly, or plausibly, related to reproductive biology, such as courtship behaviour, sex discrimination, accessory gland proteins and memory formation.

### Association between highly divergent regions and deleterious variants

To test whether highly divergent genomic regions between populations under different selection regimes could be explained by the purging of deleterious variants, we characterised the SNPs within BayPass C2 outlier regions. Of 442 C2 outlier SNPs, 90 fell within coding sequences: 81 were synonymous, one was a nonsense variant on the X chromosome, and eight were missense, of which two were confidently predicted to be deleterious by SIFT (figure S6).

We then tested whether these regions were enriched for deleterious variants that were purged under polyandry. Across 10,000 permutations, the observed number of purged deleterious variants overlapping outlier regions (2 variants) was lower than expected under the null distribution generated from synonymous variants (mean = 6.16, SD = 2.43). This deviation was not statistically significant (empirical p = 0.986, z = –1.71), indicating that highly divergent regions do not preferentially contain deleterious variants purged under polyandry. These results suggest that purifying selection against deleterious alleles is unlikely to be the primary driver of divergence in these regions.

## Discussion

Our results provide genomic evidence consistent with sexual selection enhancing the purging of deleterious alleles. Populations that evolved under strong sexual selection carried significantly fewer predicted deleterious variants, indicating reduced mutation load, than those where sexual selection was experimentally removed. This effect was consistent across both individual and population-level analyses. These findings constitute the strongest and most comprehensive empirical support to date for theoretical predictions, and prior experimental observations, that sexual selection can facilitate the removal of deleterious mutations (Rowe & Houle 1996; Whitlock & Agrawal 2009).

The observed reduction in deleterious variants under polyandry indicates that sexual selection can act as a powerful source of purifying selection. The importance of this finding is emphasised by the lack of difference observed in nucleotide diversity (π) or runs of homozygosity (F_ROH_) between treatments. This suggests that lower mutation load in sexually selected populations was not due to demographic factors, such as reduced effective population size (N_e_) linked to the treatments. Although the experimental design equalised the census-based effective population size between sexual selection regimes, realised N_e_ could still have been reduced under polyandry due to variance in male mating success. However, the absence of reduced π or increased F_ROH_ implies that any skew did not substantially alter genome-wide N_e_. This pattern suggests that reduction in deleterious alleles in the sexual selection treatment was not a by-product of demographic contraction or mating skew, but rather the outcome of more efficient selection, removing deleterious genetic variation *per se*. Under polyandry, both male–male competition and female choice likely increased the strength of selection against high-load males, reducing transmission of deleterious alleles without eroding neutral diversity. The absence of N_e_ signatures therefore supports a model in which sexual selection enhances the efficacy of selection by favouring genetically higher-quality males, rather than one in which purging arises from extreme reproductive skew.

The magnitude of purging scaled with the predicted functional impact of the variants assessed. Nonsense variants showed the strongest reduction under polyandry, followed by missense deleterious variants, while missense tolerated variants were largely unaffected. This pattern is consistent with the prediction that sexual selection should be most effective against deleterious alleles with large fitness effects if mating success is sensitive to overall condition (Rowe and Houle 1996; Whitlock and Agrawal 2009). This is the first time that such a pattern has been reported, providing the strongest genomic support to date for this hypothesis. Nonsense mutations, which often lead to a complete loss-of-function, are likely highly deleterious and therefore more efficiently removed from the population when sexual selection is strong. Missense deleterious variants, which tend to have more moderate effects, were also reduced under polyandry, but to a lesser extent, and their frequency patterns approached those of missense tolerated variants. Essentially, sexual selection acts as a genome-wide filter that preferentially removes the most damaging coding variants, rather than uniformly reducing diversity.

Despite sexual selection acting primarily through males in our experiment, we found that the females we sequenced carried fewer nonsense mutations than the males within the same treatment. This sex difference cannot be explained by overall purging of deleterious alleles, as both sexes share the same gene pool, but may instead reflect sex-specific selection pressures or mutational processes. For example, females with high mutational load may experience stronger viability selection prior to sampling meaning that only those with lower load would reach adulthood. Recent work has shown that deleterious mutations can have sex specific larval viability effects and that females are more sensitive that males to early life stress (Teder & Kaasik 2023; Melde et al. 2024). Further work is required to assess the possibility of such mechanisms in this system. However, that mutation load of both sexes is reduced under polyandry does support the notion that sexual selection enhances purging primarily through male reproductive filtering, indirectly benefiting females and the population as a whole.

Our analysis of population persistence under inbreeding further supports the conclusion that sexual selection reduces mutation load in a way that enhances population viability. This relationship parallels the earlier phenotypic findings of Lumley et al. (2015), but our results now provide direct genomic evidence that populations evolving under strong sexual selection carry significantly fewer deleterious variants. Crucially, extinction risk was best predicted by mutation load, rather than sexual selection treatment itself, consistent with mutation load being the mechanistic link between sexual selection and extinction resilience. Remarkably, the genomes sampled from the experimental lines 102 generations after the experiment performed by Lumley et al. (gens 54 vs 156) still predicted past survival implying that that differences in load between populations were both substantial and persistent. This temporal persistence may suggest that the majority of purging took place early in the experimental evolution, prior to generation 54, but this is not possible to confirm with the samples/data available.

The findings from our study contribute to a growing recognition that sexual selection is important not only in evolutionary terms but also from a conservation perspective (Cally et al. 2019). By reducing mutation load while maintaining genomic diversity, sexual selection can enhance the resilience of small or inbred populations, potentially contributing to evolutionary rescue. Indeed, the purging of mutation load through sexual selection may contribute to resolving the sex paradox, explaining the long-term maintenance of sex in nature, despite it’s inherent costs (Maynard-Smith 1978). However, sexual selection also increases variance in reproductive success and so, in some situations, may exacerbate demographic stochasticity in small populations. Whether sexual selection ultimately helps or harms threatened populations will depend on the balance between its genetic benefits and demographic costs, which warrants future theoretical and empirical work.

In our study, genome scans revealed multiple peaks of divergence between populations evolved under differential strengths of sexual selection. The genes located in these divergent regions had diverse functions, including protein synthesis, lipid metabolism, and oxidative stress tolerance, as well as several that were directly linked to reproduction—such as courtship behaviour, sex discrimination, accessory gland proteins, and memory formation (Gillott 2003; Scheunemann et al. 2019; Sellami et al. 2015). These functions align with those seen to evolve in response to sexual selection in other systems (e.g. Wiberg et al. 2021; Veltsos et al. 2022; Wyer et al. 2023; Fromonteil et al. 2025). The diversity of functions suggests that experimental sexual selection influences a broad suite of physiological and behavioural traits, consistent with the polygenic nature of reproductive fitness, but focused on facets of reproductive biology key to both pre- and post-mating reproductive success (recognition, courtship behaviour as well as seminal fluid protein functions).

Importantly, divergent genomic regions were not enriched for deleterious variants that were at lower frequency in polyandrous populations. This finding suggests that genomic divergence between the polyandrous and monandrous populations is not the result of differential purging, but instead reflects positive selection for beneficial alleles, especially those associated with reproductive success.

Our findings are consistent with the predictions of the “good genes” model, which proposes that sexual selection favours individuals carrying alleles that enhance fitness. Under the ‘genic capture’ good genes model, sexually selected traits are condition-dependent and reflect the cumulative effects of many loci (Rowe and Houle 1996, Tomkins et al. 2004). By capturing genetic variation across many loci affecting condition, these traits allow sexual selection to continually act without depleting the underlying variation (and thus potentially explain the lek paradox. In our *Tribolium* populations, polyandrous lines carried significantly fewer high-impact deleterious variants—particularly nonsense and deleterious missense alleles—indicating efficient genome-wide purging. At the same time, divergent genomic regions were generated by the experimental evolution that were enriched for genes associated with reproductive functions but not for deleterious variants, suggesting that adaptive divergence occurs largely independently of purging.

Together, our results show that sexual selection can operate via two complementary processes: enhancing purifying selection on deleterious alleles across the genome while simultaneously driving adaptive differentiation at loci underpinning reproductive success. By linking functionally annotated mutation load to extinction risk in experimentally evolved populations, our results provide direct evidence of the genomic mechanisms by which sexual selection promotes both genomic health and adaptive divergence, thereby helping to maintain sexual reproduction and enhance population resilience.

In providing evidence that sexual selection reduces mutation load and increases population viability, our study supports the possibility that this mechanism may help explain the widespread prevalence of sexual reproduction in nature, despite the costs of such reproduction (Maynard-Smith 1978). The findings also highlight the importance of allowing sexual selection to act in sexually reproducing populations of conservation concern, given its potential to reduce mutation load without reducing overall genomic diversity.

## Materials and Methods

### Beetles and husbandry

Beetles were of the Georgia-1 (GA1) strain, collected from the wild in 1980 and kept by the Beeman laboratory at the US Department of Agriculture. Once acquired by our laboratory, the stock population was maintained on a standard medium of 90% organic wheat flour and 10% brewer’s yeast, which we refer to as ‘fodder’, at 30°C, 60% relative humidity, and a 12:12 light-dark cycle (light from 0800-2000). The standard husbandry cycle consisted of two phases; during the oviposition phase, adult (12+/−3 days post-eclosion) beetles randomly chosen to parent the next generation were removed from their populations and placed into fresh fodder for seven days of mating and egg-laying. Following this period, adults were sieved from the fodder and discarded, beginning a 35-day development phase—during which time, eggs in the fodder developed through larval and pupal stages to become adults. By preventing any interaction between sexually mature adults and offspring, this regime of non-overlapping generations reduced the risk of negative density-dependent effects, eliminated opportunities for intergenerational interactions such as egg-cannibalism, and allowed accurate tracking of generations.

### Experimental sexual selection regimes

As described in Lumley et al. (2015), unmated adult beetles from a single GA1 stock population (maintained at a size of 600 adults) were randomly assigned to begin each sexual selection regime. By manipulating the number of males and females housed together during the oviposition phase, differential levels of sexual selection were imposed on each regime while controlling for population size. The experiment consisted of six lines, representing three independent replicates to two selection regimes. Within each of three independent monandrous lines, the eggs produced by 20 independent single-pair matings were combined prior to the development phase. In three independent polyandrous lines, each of 12 females was housed with five males, with the eggs produced being combined for development. Thus, monandry removed all opportunity for female choice and male-male competition, whereas polyandry imposed sexual selection through competition between males for a single female, and provided that female with a choice between five males. The numbers of pairs used for monandry and groups used for polyandry were defined in order to equalise the effective population size, N_e_, for mixed adult sex ratios 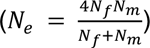 across treatments (Falconer 1996; see also Lumley et al. 2015). Thus *N_e_* = 40 in both regimes. Treatments were applied for 156 beetle generations (15 years) prior to the commencement of this study.

### Sample preparation and sequencing

Unmated adult beetles were collected from each replicate of the two sexual selection regimes, and flash-frozen in liquid nitrogen. For each individual DNA extraction was conducted using the DNeasy blood and tissue kit (insect tissue protocol, Qiagen), with the whole individual ground in liquid nitrogen. The extract was purified using a 1x bead cleanup with the AMPure XP SPRI protocol (Beckman Coulter). Library preparation and sequencing were performed at the Earlham Institute (Norwich, UK) using the low-input transposase-enabled (LITE) pipeline (see supplementary methods). Samples were sequenced on two flowcells across two sequencing runs, the first on the Illumina Novaseq 6000 and the second on the NovaSeq X Plus platform. Samples from all treatments were randomly distributed across both runs to avoid confounding treatment effects with sequencing platform or run-specific variation. Sequences were obtained from 84 individuals: 42 from each sexual selection regime, divided equally between each replicate line and between the sexes.

### Variant calling and filtering

Reads were trimmed using Trimmomatic v0.39 (Bolger, Lohse and Usadel, 2014) and mapped to the Tcas5.2 reference genome (Herndon et al 2020) using BWA-MEM v0.7.17 (Li, 2013). Mapping was followed by SAMtools v1.18 *fixmate* and *sort* to ensure BAM files were correctly formatted and sorted prior to removing duplicates (Danecek et al., 2021). PCR duplicates were removed using Picard v2.26.2 *MarkDuplicates* (Broad Institute, 2019). Finally, mappings were filtered for complete read pairs with a mapping quality (MAPQ) >25 using SAMtools *view*. Joint genotyping was conducted using BCFtools v1.18.0 mpileup (Danecek et al., 2021), with ploidy specified by sex and chromosome to ensure haploid genotyping of the X chromosome in males. BCFtools *call* was then used to call all sites under the multi-allelic model (-m). BCFtools *filter* was used to remove variants within 3bp of other variants, with a variant quality score <30, that were at a locus with sequencing depth less than 578 and greater than 5201 (+/- 3 times average sequencing depth), and were represented by data at that locus in less than 50% of individuals (-g 3 -G 3 -e ‘DP < 578 || DP > 5201 || F_MISSING > 0.5 || QUAL < 30’). The resulting file is referred to hereafter as the allsites vcf.

At this point, from the allsites vcf single nucleotide polymorphisms (SNPs) were extracted using BCFtools view and further filtered to remove sites with minor allele count <3. This file is referred to hereafter as the SNP vcf. To prepare the allsites vcf for further analysis, variant and invariant sites were handled separately: Invariant sites were extracted using VCFtools 0.16.0 (Danecek et al. 2011) and stored in a separate file. Variant sites were filtered using VCFtools to only contain biallelic SNPs that did not deviate significantly from Hardy-Weinberg Equilibrium (*HWE* p-value < 0.001). Following this filter, invariant and variant sites were concatenated using BCFtools *index* and *concat*. This file is referred to hereafter as the filtered allsites vcf.

Quality control of the SNP VCF revealed three samples (polyAmT1, monoCmT, monoAfT) with huge counts of singleton SNPs and indels compared to other samples. We excluded reads from these samples from the raw sequence data and reran the above steps to regenerate both the SNP vcf and the allsites VCF. After filtering, we took forward a set of 2,874,861 SNPs for analysis.

As samples were sequenced across two runs on different Illumina platforms, we used principal component analysis (PCA) to assess whether platform- or run-specific effects contributed to structure in the data. The SNP VCF was pruned for linkage with bcftools (*+prune -m 0.3 -w 50kb*) before PCA with plink v1.9 (Purcell et al. 2007) on a set of 62,713 SNPs. Using the first and second principal components generated by this analysis, which accounted for 29.1% and 19.6% of variance, respectively, we observed no clustering of samples by sequencing run (figure S1); sequencing run was therefore not used as a covariate in subsequent analyses.

### Assessing inbreeding

As a proxy for inbreeding in experimental populations, we quantified runs of homozygosity (ROH) using plink v1.9. Analyses were restricted to autosomal linkage groups, and ROH were called using the following parameters: a minimum length of 200kb (*--homozyg-kb 200*) and at least 50 SNPs per ROH (*--homozyg-snp 50*), with a minimum SNP density of one SNP per 25kb (*--homozyg-density 25*). Gaps of <100kb were allowed between consecutive homozygous SNPs within a ROH (*--homozyg-gap 100*). A sliding window approach was applied with 50 SNPs per window (*--homozyg-window-snp 50*), allowing up to 1 heterozygous call *(--homozyg-window-het 1*) and 3 missing calls *(--homozyg-window-missing 3*) per window. A window was required to be called homozygous in at least 5% of its positions to contribute to an ROH *(--homozyg-window-threshold 0.05*). These parameters largely follow PLINK defaults and those used in analyses of insects and other organisms.

The genomic inbreeding coefficient (F_ROH_) was computed, per individual, as the proportion of that individual’s genome in ROH (‘KB’ field returned by plink divided by the combined callable length of all autosomes). Differences in F_ROH_ between treatments were assessed using the non-parametric Wilcoxon rank-sum test, as the data were right-skewed and could not be transformed to fit a normal distribution.

### Estimating mutation load

SNPs within coding sequences (CDS) of the Tcas5.2 annotation were extracted and functionally annotated using SnpEff v4.3 (Cingolani et al. 2012), which assigned each variant to an effect category (nonsense, missense, or synonymous). Missense variants were further categorised using SIFT 4G (Vaser et al. 2016), with the provided *Tribolium castaneum* database, to predict the strength of their potential functional impact. SIFT uses a database of known proteins to score the putative effect of a change of amino acid at each position in the protein based on the frequency of that substitution in a multi-species alignment, and the biochemical similarity of the amino acids. Each SNP obtains a SIFT score in the range 0-1, with values of <0.05 considered deleterious, and an estimate of confidence in the assigned score. In using SIFT annotations we excluded any sites tagged as low confidence.

The resulting annotation allowed coding SNPs to be divided into four categories of predicted functional impact: a) synonymous, causing no change in the coded amino acid; b) missense tolerated, amino acid substitutions predicted not to impact protein function based on patterns of protein conservation; c) missense deleterious, amino acid substitutions predicted to disrupt protein function based on patterns of protein conservation; d) nonsense, the introduction of premature stop codons. Per-individual counts of SNPs in each category were used as a relative metric of mutation load, avoiding the need to make assumptions about the average dominance, which would be required in attempting to translate variant counts into explicit estimates of fitness effects. Differences in individual sequencing coverage and overall genetic diversity across populations were accounted for by calculating, per individual, the ratio of variants in a focal category to the number of synonymous variants observed in the same individual.

Separately for autosomal nonsense, missense deleterious and missense tolerated variant categories, we fitted GLMMs (lme4 with additional statistics from lmerTest; Kuznetsova et al. 2017) in which the response variable was the ratio of the number of variants of the focal category/the number of synonymous variants, per individual. Initial models included sexual selection regime, sex and their interaction as fixed effects, with the replicate line ID and individual ID as nested random factors. Interactions were removed if non-significant. Pairwise comparisons were performed post-hoc using emmeans (Lenth & Piakowski 2025). Models of the same structure were fitted to test the same variant categories on the X chromosome.

We also evaluated mutation load at the population level. For this analysis, individuals from replicate lines within a treatment were pooled and treated as a single treatment-level population. Using only coding SNPs, for each category of variant impact, at each site *i* we computed the observed allele frequency of the alternative allele in a population X as f^x^_i_ = d^X^_i_ / n^X^_i_, where n^X^_i_ is the total number of alleles called at *i* in population X and d^X^_i_ is the number of alternative alleles called. We computed R_xy_ (described by Xue et al. 2015; after the method of Dussex et al. 2023) for each functional category between monandrous and polyandrous populations, to assess the relative frequency of deleterious variants under each sexual selection regime. Analyses were restricted to sites that were polymorphic in at least one of the compared populations, thereby capturing variants that may have been completely purged from one group. Each R_xy_ value was standardised using the same statistic computed on a random subset of intergenic variants from the same populations. The variance in R_xy_ was estimated by block jackknifing across the data, systematically removing non-overlapping blocks of SNPs 10% of the size of the data (Dussex et al. 2023).

### The effect of total load on survival under inbreeding

We reanalysed survival data collected from generation 54 of our experimental populations by Lumley et al. (2015) to test whether the differential survival under inbreeding observed in this study is more closely associated with our estimate of mutation load or with the original line-level classification of sexual selection regime. These data track the survival of lineages through multiple generations of enforced sib x sib inbreeding (details in Lumley et al. 2015). We fitted Cox proportional hazards mixed effects models (coxme package) to survival data from Lumley et al., where survival time was defined as the number of generations to extinction, with surviving populations treated as right-censored. Three models were fitted, each with survival time as the dependent variable, and replicate line within the selection regime as a random factor. In each case, the proportional hazards assumption was verified with the cox.ph function. Independent fixed factors in each model were: 1) sexual selection regime, 2) mutation load proxied by counts of deleterious sites 3) sexual selection regime + mutation load.

Using our mutation load estimate, we calculated a metric representing the total effect of deleterious variants that are on average partially recessive (Huber et al. 2018; Kyriazis & Lohmueller 2024), where variants predicted as nonsense are assigned as ‘more deleterious’ than those predicted to be missense deleterious (as indicated by our mutation load analysis, see results):

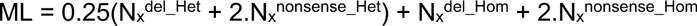

Where N_x_ represents the ratio of the number of variants of a focal category to the number of synonymous variants observed in the same individual, *del* and *nonsense* refer to missense deleterious and nonsense categories of variants, and *Het* and *Hom* refer to variants in heterozygous and homozygous form respectively. To test the sensitivity of results to this parameter, we repeated the analysis across nine combinations of mean dominance and relative weighting of missense deleterious and nonsense variant effects (see supplementary methods). Results were qualitatively equivalent across load metrics. We therefore present the original case here with further detail provided in the supplementary material.

### Regions of strong divergence

We performed a genome-wide scan for selection using BayPass v2.4 (Gautier 2015), implementing a Bayesian framework sensitive to demography. BayPass estimates a background allele frequency (omega) matrix across populations, to account for the confounding effect of demography which can frustrate the identification of selected variants (Günther and Coop 2013; Gautier 2015). This approach allowed us to control for unquantified differences in relatedness among individuals used to found each selection line. The BayPass model computes an omega matrix to correct for neutral correlations when testing allele frequencies for population divergence or association with environmental/trait variables. Within BayPass, we utilised the statistic C2, which contrasts SNP allele frequencies between two groups of populations specified by a binary trait. This method outperforms others in identifying SNPs under selection in scenarios analogous to our experiment (Olazcuaga et al. 2020).

We computed C2 across the two sets of 3 replicate sexual selection lines, with sexual selection regime as the binary covariable. To avoid the impact of small, annotation-sparse, unplaced scaffolds in the reference genome, we ran BayPass on the 10 linkage-group-level scaffolds. We pre-computed the BayPass omega matrix using a curated subset of independent, high confidence, highly representative, putatively neutral SNPs which afforded the best opportunity to estimate the neutral covariance in allele frequencies across the genome (omega dataset; see supplementary methods). BayPass was run independently for autosomal and X-linked loci to reflect their different modes of inheritance. For autosomal runs, the omega dataset consisted of 7,158 non-exonic SNPs, visually confirmed to be evenly spread across the genome. Omega dataset for the X-chromosome consisted of 820 SNPs. The dataset used in autosomal BayPass analysis runs contained a less stringently filtered set of 1,833,626 sites, derived from the SNP vcf (see Supplementary Methods; 26,259 for the x-linked analysis run).

For each analysis, we performed two independent BayPass runs with different random seed initiators and computed correlations to test the consistency of model performance with our data (Olazcuaga et al. 2020; Dickson et al. 2020). The C2 estimates were calibrated using a pseudo-observed dataset (POD; Gautier 2015; see supplementary methods). The 0.999 quantile of C2 values from the POD analysis was used as the outlier threshold above which to identify C2 candidate SNPs. C2 candidate regions were defined as those containing >=2 outlier SNPs separated by <50kb (Gautier 2015), with top candidate SNPs being those in each candidate region with the highest value of C2.

### Effects of BayPass outliers

To assess the effects of BayPass outliers on phenotype, we collected functional annotations of coding genes in proximity to these loci. First, we concatenated BayPass outliers identified on the autosomal and X linkage groups, then derived two lists of variants of interest: a) all SNPs with significant C2 (C2 candidate SNPs), to examine divergent regions; b) the SNP with highest C2 from each candidate region (top candidate SNPs), to examine sites more likely to be driving divergence. From each of these lists, a gene set was generated by intersecting their positions with the genome annotation (Tribolium_castaneum.T.cas5.2.59.gff3), one containing the closest gene to each C2 candidate (bedtools -*closest*), and one containing all genes within 10kb of a top candidate SNP (bedtools -*slop*, *-intersect*). We used these gene lists as input to g:Profiler (Reimand *et al*. 2007; Kolberg *et al*. 2023) to test for enrichment of functional terms derived from gene ontology (GO). All other settings were the G:profiler defaults, the background used was all genes in the *T. castaneum* genome. The OSG3 annotation (https://ibeetle-base.uni-goettingen.de/download/species/Tcas/OGS3.gff.gz) and the iBeetle-Base database (Dönitz *et al*. 2018) were used to manually identify gene functions, and orthologous genes in *Drosophila* melanogaster were subsequently characterised using Flybase (Öztürk-Çolak *et al*. 2024).

### Association between highly divergent regions and deleterious variants

To test whether divergence in candidate regions was driven by purging of deleterious variants, we asked whether highly divergent genomic regions (BayPass C2 outliers) were enriched or depleted for variants predicted to be deleterious (missense deleterious or nonsense) and purged under polyandry (i.e., observed at lower frequency under polyandry relative to monandry). Only outlier regions overlapping coding sequences were included, as some fell entirely in non-coding regions and could not contain annotated variants. For each deleterious variant subset, we compared the observed overlap with a null expectation generated by randomly sampling an equal number of synonymous variants 10,000 times. For each permutation, we counted the number of sampled variants overlapping the outlier regions, producing a null distribution of expected overlap. We calculated an empirical p-value as the proportion of permutations in which the overlap equalled or exceeded the observed value, and a z-score to quantify deviation from the null expectation.

## Acknowledgments

We thank Alyson Lumley who worked on founding the lines used here, along with all members of the Tribolium lab at UEA who contributed time and effort to their maintenance. We thank Maria-Elena Mannarelli for wetlab work on the samples.

The work was funded by Natural Environment Research Council grant NE/T007885/1. WN acknowledges support from the Biotechnology and Biological Sciences Research Council (BBSRC), part of UK Research and Innovation, Core Capability Grant BB/CCG2220/1 at the Earlham Institute (EI) and its constituent work packages (BBS/E/T/000PR9818 and BBS/E/T/000PR9819), and the Core Capability Grant BB/CCG1720/1 and the National Capability BBS/E/T/000PR9816 (NC1—Supporting EI’s ISPs and the UK Community with Genomics and Single Cell Analysis), BBS/E/T/000PR9811 (NC4—Enabling and Advancing Life Scientists in data-driven research through Advanced Genomics and Computational Training), and BBS/E/T/000PR9814 (NC 3—Development and deployment of versatile digital platforms for ‘omics-based data sharing and analysis). Also support from BBSRC Core Capability Grant BB/CCG1720/1 and the work delivered via the Scientific Computing group, and the physical HPC infrastructure and data centre delivered via the NBI Computing infrastructure for Science (CiS) group. For the purpose of open access, the author has applied a Creative Commons Attribution (CC BY) licence to any Author Accepted Manuscript version arising from this submission.

