## Supplementary_material_pdf for "Sexual selection purges mutation load, but not overall genetic diversity in populations, decreasing vulnerability to extinction"

### Supplementary methods

#### The effect of mutation load on survival under inbreeding

Time to extinction under inbreeding was modelled with a range of different metrics of mutation load, corresponding to different assumptions about the mean dominance and relative weighting of missense deleterious and nonsense variant effects (table S1).

**Table S1.** Combinations of dominance and weighting of missense deleterious:nonsense variant effects used to compute nine metrics of mutation load.

| | Mean dominance of alt allele (h) | | |
|---|---|---|---|
| | Totally recessive (h = 0 | Partially recessive (h = 0.25) | Codominant (h = 0.5) |
| Relative weight (nonsense / missense deleterious; w) | w = 1, h = 0 | w = 1, h = 0.25 | w = 1, h = 0.5 |
| | w = 2, h = 0 | w = 2, h = 0.25 | w = 2, h = 0.5 |
| | w = 5, h = 0 | w = 5, h = 0.25 | w = 5, h = 0.5 |

The metric of mutation load in each case was computed as:

$$ML = h(N_x^{del\_Het} + w.N_x^{nonsense\_Het}) + N_x^{del\_Hom} + w.N_x^{nonsense\_Hom}$$

Where $N_x$ represents the ratio of the number of variants of a focal category to the number of synonymous variants observed in the same individual, *del* and *nonsense* refer to missense deleterious and nonsense categories of variants, and *Het* and *Hom* refer to variants in heterozygous and homozygous form respectively.

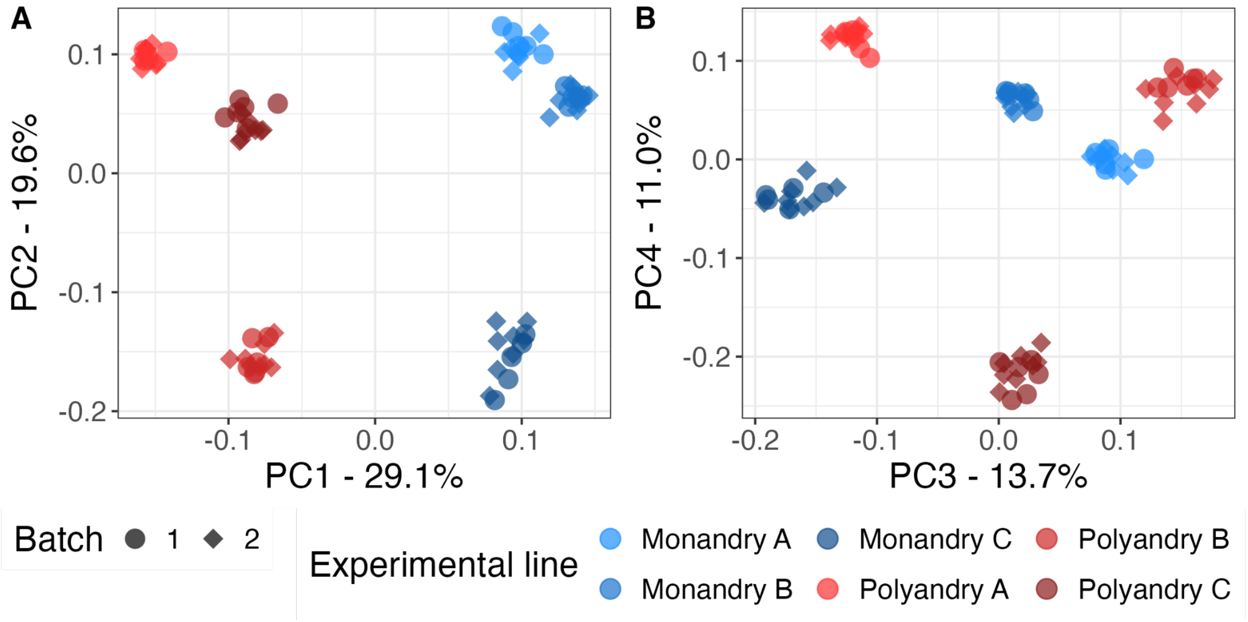

**Figure S1. No batch effect from sequencing runs is evident in the data**
Principle components summarising genomic variation of *T. castaneum* populations (colour shade) experimentally evolved under monandrous and polyandrous sexual selection regimes (blue/red). The shape of points indicates in which of two batches the samples were sequenced. No batch effect is evident. The most explanatory principle component (PC1) appears to capture the difference between sexual selection regimes.

### Supplementary results

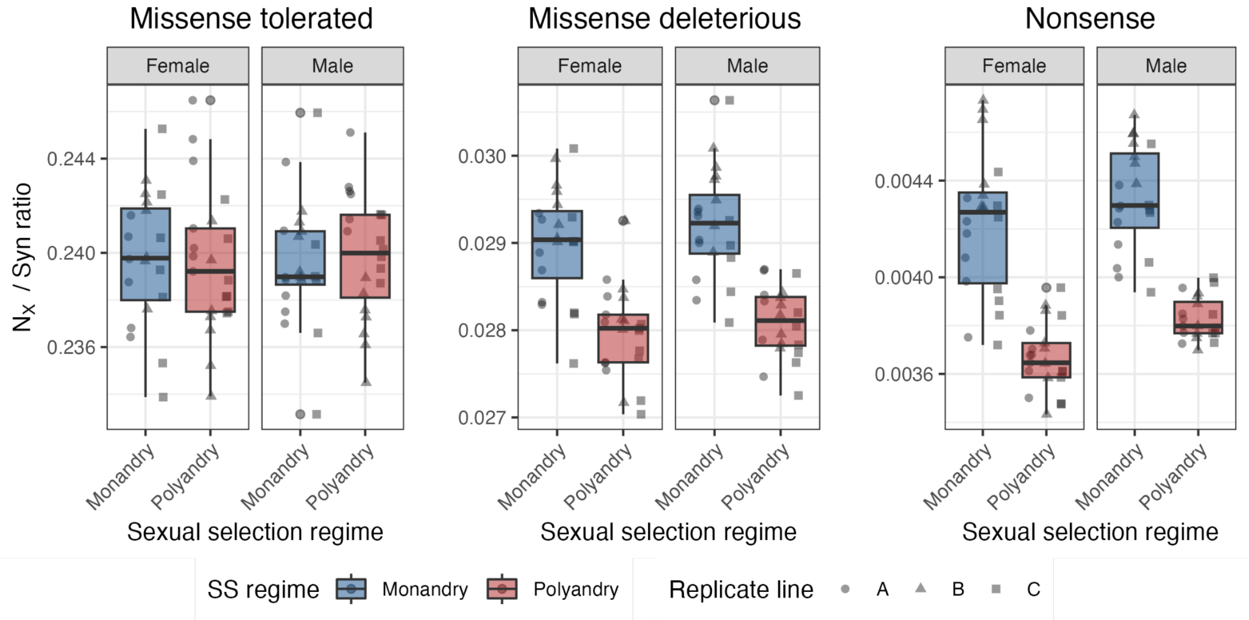

**Figure S2. Females carry a higher burden of deleterious variants than males**
The effect of sexual selection on individual mutation load of males and females on the autosomes. The ratio of the number missense tolerated, missense deleterious and nonsense ($N_X$) variants to the number of synonymous variants per individual, for populations of *Tribolium castaneum* evolving under either monandrous or polyandrous sexual selection regimes for 156 generations. Plots for each functional category are stratified by sex.

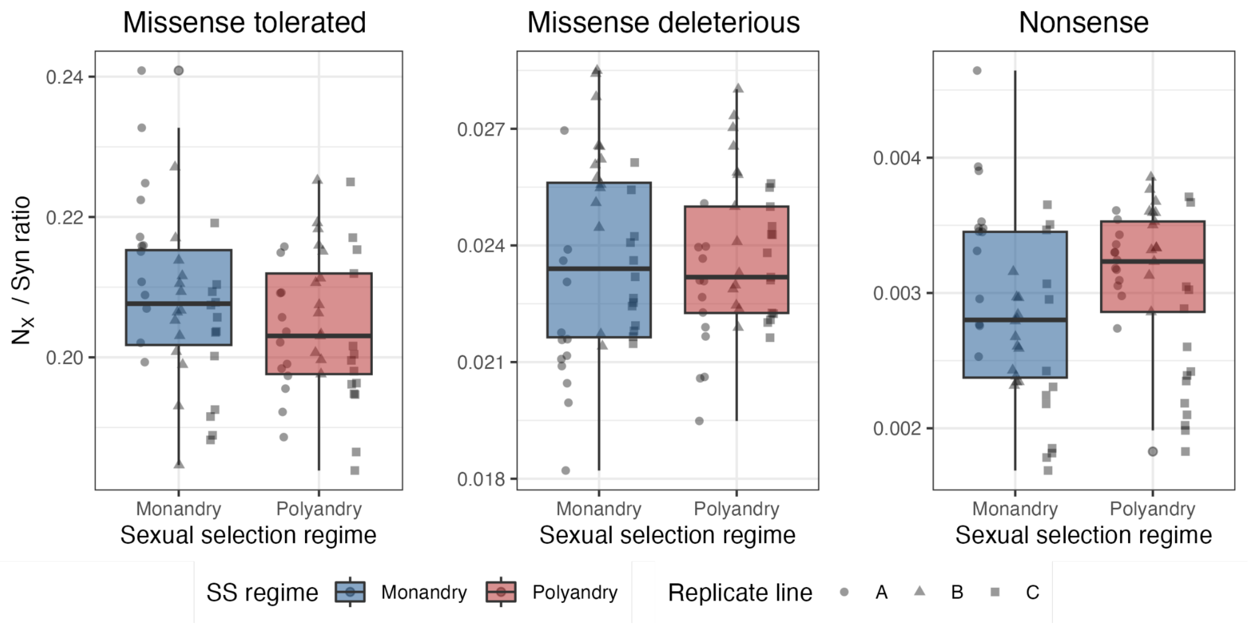

**Figure S3. Mutation load on the X chromosome.**
Genetic variation on the X chromosome categorised by increasing deleterious functional impact (misense tolerated - nonsense) in populations of Tribolium castaneum evolved under either monandrous or polyandrous sexual selection regimes for 156 generations. Points show the ratio of missense tolerated, missense deleterious and nonsense (NX) sites to synonymous sites, per individual.

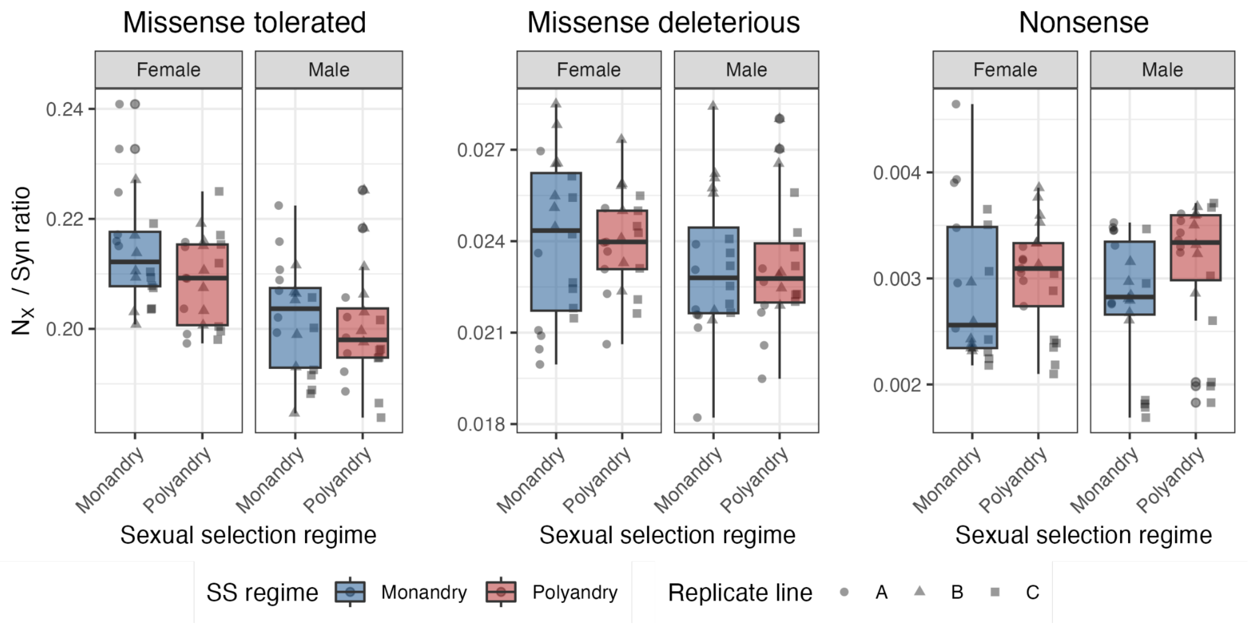

**Figure S4. Mutation load on the X chromosome by sex.**
The effect of sexual selection on individual mutation load of males and females on the X chromosome. The ratio of the number missense tolerated, missense deleterious and nonsense (NX) variants to the number of synonymous variants per individual, for populations of Tribolium castaneum evolving under either monandrous or polyandrous sexual selection regimes for 156 generations. Plots for each functional category are stratified by sex.

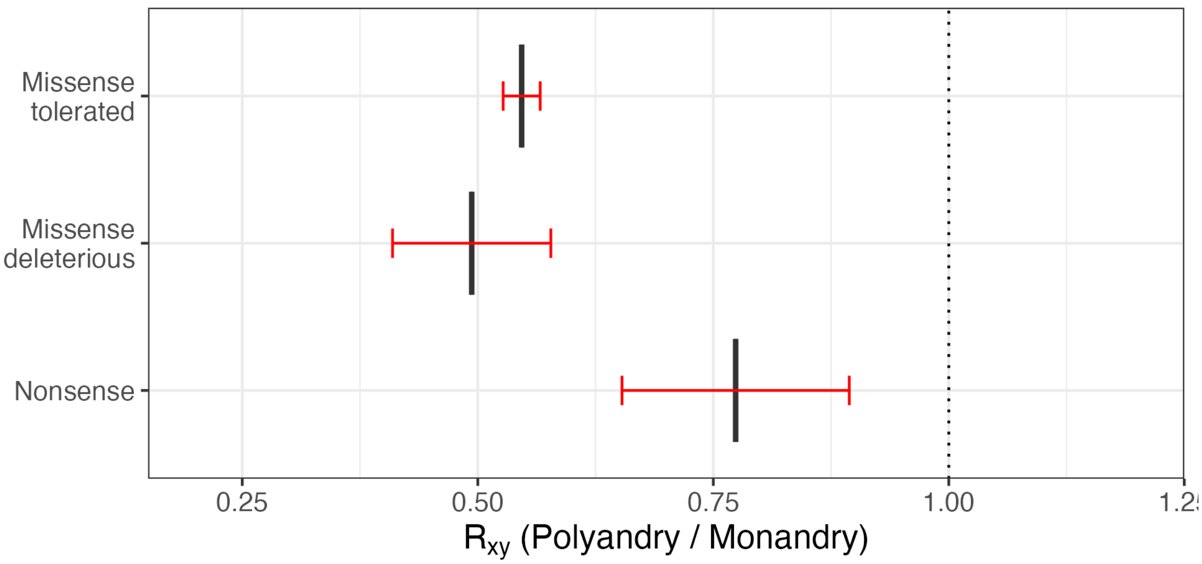

**Figure S5. The pattern of purging observed on the autosomes is not evident on the X chromosome.**
Genetic variation categorised by increasing deleterious functional impact in populations of Tribolium castaneum evolved under either monandrous or polyandrous sexual selection regimes for 156 generations. $R_{xy}$ analysis contrasting allele frequencies in monandrous and polyandrous populations, across functional categories of high-quality SNPs, normalized by Rxy of intergenic variants. $R_{xy} < 1$ indicates a relative frequency deficit of the corresponding category under polyandry. Red bars represent 95% CIs computed from block jack-knife standard errors.

**Mutation load and survival under inbreeding**

We compared ability of SS regime and mutation load to predict extinction risk. The main text presents results from the scenario we thought most likely to represent biological reality (core scenario), but to assess the robustness of the result to the load metric used, we repeated the analysis a total of nine metrics of mutation load representing different assumptions about the mean dominance of variants (h) and the relative mean strengths of nonsense variants to missense deleterious and variant effects (w; see supplementary methods).

In the core scenario, extinction time was best predicted by mutation load, with the mutation load model having an AIC at least two less than the SS regime model. This same pattern was observed across every mutation load metric tested, except when $h = 0$ & $w = 5$, corresponding to a scenario where deleterious variants are totally recessive and nonsense variants are on average 5 times more deleterious than missense deleterious ones (table S2).
In models fitting both SS regime and mutation load together as fixed factors, SS regime was never a significant predictor of time to extinction. Mutation load remained marginally significant <0.069) in eight of the nine models but its significance dropped relative to being fit alone. This is expected given that we know SS regime and mutation load to be highly colinear.

Overall, the result that mutation load best explained extinction risk was extremely robust to alternative methods of converting counts of variants to estimates of realised mutation load. Full model outputs and fit comparisons are given in table S2.

**Table S2.** Functional characterisation of genomic variants was used as an estimate of mutation load in 84 genomes sequenced from *Tribolium* castaneum lines evolved under strong (polyandry) or weak (monandry) sexual selection (SS) regimes for 156 generations. Mixed models (GLMM) were fitted to population survival data, when families derived from these same lines where subjected to enforced inbreeding (from Lumley et al. 2015), modelling survival as a product of mutation load or SS regime or both. The main text shows that the mutation load estimate alone was the best predictor of survival. To evaluate the sensitivity of this result to assumptions, we computed the metric in nine different ways, corresponding to different assumptions about the mean dominance and relative weighting of missense deleterious and nonsense variant effects. **A)** The agreement between models using different metrics of mutation load. **B)** Model comparison using AIC.

**A**

| Load metric | Model | SS regime effect (coef) | Mutation load effect (coef) | Random effect (rep: sd) |
|---|---|---|---|---|
| na | SS regime | -0.730, p = 0.026 | – | 0.020 |
| h = 0, w = 1 | Mutation load | – | 719.3, p = 0.005 | 0.009 |
| | SS regime + mutation load | -0.035, p = 0.944 | 699.0, p = 0.069 | 0.009 |
| h = 0.25, w = 1 | Mutation load | – | 378.9, p = 0.003 | 0.009 |
| | SS regime + mutation load | -0.002, p = 0.997 | 378.4, p = 0.042 | 0.009 |
| h = 0.5, w = 1 | Mutation load | – | 236.2, p = 0.844 | 0.009 |
| | SS regime + mutation load | -0.092, p = 0.844 | 220.4, p = 0.051 | 0.009 |
| h = 0, w = 2 | Mutation load | – | 632.6, p = 0.006 | 0.009 |
| | SS regime + mutation load | 0.376, p = 0.56 | 862.8, p = 0.061 | 0.009 |

| | | | | |
|---|---|---|---|---|
| h = 0.25, w = 2 | Mutation load | – | 299.5, p = 0.0029 | 0.009 |
| | SS regime + mutation load | 0.065, p = 0.900 | 314.8, p = 0.046 | 0.009 |
| h = 0.5, w = 2 | Mutation load | – | 183.3, p = 0.003 | 0.009 |
| | SS regime + mutation load | -0.071, p = 0.88 | 173.6, p = 0.054 | 0.009 |
| h = 0, w = 5 | Mutation load | – | 331.8, p = 0.020 | 0.011 |
| | SS regime + mutation load | 0.139, p = 0.906 | 390.1, p = 0.449 | 0.011 |
| h = 0.25, w = 5 | Mutation load | – | 109.1, p = 0.003 | 0.009 |
| | SS regime + mutation load | -0.047, p = 0.925 | 105.2, p = 0.059 | 0.009 |
| h = 0.5, w = 5 | Mutation load | – | 109.1, p = 0.003 | 0.009 |
| | SS regime + mutation load | -0.047, p = 0.925 | 1.05.2, p = 0.059 | 0.009 |

**B**

| **Load metric** | **Model** | **LokLik** | **AIC** | **ΔAIC from best model** | **AIC weight** |
|---|---|---|---|---|---|
| h = 0, w = 1 | SS regime | -138.1 | 280.2 | 3.13 | 0.132 |
| | Mutation load | -136.5 | 277.1 | – | 0.634 |
| | SS regime + mutation load | -136.5 | 279.1 | 2.00 | 0.234 |
| h = 0.25, w = 1 | SS regime | -138.1 | 280.2 | 4.071 | 0.087 |
| | Mutation load | -136.1 | 276.1 | – | 0.667 |
| | SS regime + mutation load | -136.1 | 278.1 | 2.00 | 0.245 |
| | SS regime | -138.1 | 280.2 | 3.722 | 0.102 |

| | | | | | |
|---|---|---|---|---|---|
| h = 0.5, w = 1 | Mutation load | -136.2 | 276.5 | – | 0.653 |
| | SS regime + mutation load | -136.2 | 278.4 | 1.961 | 0.245 |
| h = 0, w = 2 | SS regime | -138.1 | 280.2 | 4.950 | 0.055 |
| | Mutation load | -136.6 | 275.3 | – | 0.661 |
| | SS regime + mutation load | -136.5 | 276.9 | 1.675 | 0.285 |
| h = 0.25, w = 2 | SS regime | -138.0 | 280.0 | 4.07 | 0.087 |
| | Mutation load | -136.0 | 276.0 | – | 0.667 |
| | SS regime + mutation load | -136.0 | 278.0 | 2.00 | 0.245 |
| h = 0.5, w = 2 | SS regime | -138.1 | 280.2 | 3.898 | 0.094 |
| | Mutation load | -136.3 | 276.3 | – | 0.661 |
| | SS regime + mutation load | -136.3 | 278.3 | 1.984 | 0.245 |
| h = 0, w = 5 | SS regime | -138.1 | 280.2 | 0.566 | 0.355 |
| | Mutation load | -137.8 | 278.6 | – | 0.471 |
| | SS regime + mutation load | -137.8 | 281.6 | 1.986 | 0.174 |
| h = 0.25, w = 5 | SS regime | -138.1 | 280.2 | 3.490 | 0.113 |
| | Mutation load | -136.4 | 276.7 | – | 0.648 |
| | SS regime + mutation load | -136.4 | 278.7 | 1.991 | 0.239 |
| h = 0.5, w = 5 | SS regime | -138.1 | 280.2 | 3.490 | 0.113 |
| | Mutation load | -136.4 | 276.7 | – | 0.648 |
| | SS regime + mutation load | -136.4 | 278.7 | 1.991 | 0.239 |

**Table S4. Bypass outlier SNPs.**

The 442 SNPs identified as significantly associated with sexual selection regime using BayPass. The closest gene to each SNP, the distance and the genomic feature and gene description were obtained by intersecting SNP positions with the genome annotation (Tribolium_castaneum.Tcas5.2.59.gff3). Drosophila homologs were identified with ibeetlebase and characterised with FlyBase.

| Chrom | SNP pos | Closest gene | Distance | Feature | Description from T.cast annotation | Fly homolog ID | Fly homolog name |
|---|---|---|---|---|---|---|---|
| LG10 | 2875218 | TC001520 | 744 | - | - | - | - |
| LG10 | 2876118 | TC001520 | 1644 | - | - | - | - |
| LG10 | 2888791 | TC001531 | 0 | intron | - | - | - |
| LG10 | 2890963 | TC001531 | 0 | intron | - | - | - |
| LG10 | 2899728 | TC001531 | 0 | intron | - | - | - |
| LG10 | 2899752 | TC001531 | 0 | intron | - | - | - |
| LG10 | 3148149 | TC034902 | 9 | intron | UPF0729 protein CG18508-like protein | FBgn0028746 | CG18508 |
| LG10 | 3148218 | TC034902 | 0 | 3` UTR | " | " | " |
| LG10 | 3148505 | TC034902 | 0 | 5' UTR | " | " | " |
| LG10 | 3148505 | TC011182 | 0 | exon | 60S ribosomal protein L7a-like Protein | FBgn0014026 | Ribosomal protein L7A |
| LG10 | 3148524 | TC034902 | 0 | 5' UTR | UPF0729 protein CG18508-like protein | FBgn0028746 | CG18508 |
| LG10 | 3148524 | TC011182 | 0 | intron | 60S ribosomal protein L7a-like Protein | FBgn0014026 | Ribosomal protein L7A |
| LG10 | 3148657 | TC011182 | 0 | intron | " | " | " |
| LG10 | 3148712 | TC011182 | 0 | intron | " | " | " |
| LG10 | 3148988 | TC011182 | 0 | exon | " | " | " |
| LG10 | 3149014 | TC011182 | 0 | exon | " | " | " |
| LG10 | 3149472 | TC011182 | 0 | intron | " | " | " |
| LG10 | 3149481 | TC011182 | 0 | intron | " | " | " |
| LG10 | 3149493 | TC011182 | 0 | intron | " | " | " |
| LG10 | 3150585 | TC011181 | 0 | exon | Serine/threonine-protein phosphatase | FBgn0003139 | Protein phosphatase V |
| LG10 | 3151854 | TC011310 | 0 | exon | Pre-mRNA-processing factor 17-like Protein | FBgn0038927 | CG6015 |
| LG10 | 3151998 | TC011310 | 0 | exon | " | " | " |
| LG10 | 3152469 | TC011310 | 0 | intron | " | " | " |

| | | | | | | | |
|---|---|---|---|---|---|---|---|
| LG10 | 3152507 | TC011310 | 0 | intron | " | " | " |
| LG10 | 3153195 | TC011310 | 0 | exon | " | " | " |
| LG2 | 6360635 | TC000829 | 0 | intron | Serine protease H18 | FBgn0035496 | Serine protease homolog (var) |
| LG2 | 6360646 | TC000829 | 0 | intron | " | " | " |
| LG2 | 6361627 | TC000829 | 0 | exon | " | " | " |
| LG2 | 6361627 | TC030253 | 0 | intron | Gustatory receptor | - | - |
| LG2 | 6361719 | TC000829 | 0 | exon | Serine protease H18 | FBgn0035496 | Serine protease homolog 35 |
| LG2 | 6361719 | TC030253 | 0 | intron | Gustatory receptor | - | - |
| LG2 | 6361730 | TC000829 | 0 | exon | Serine protease H18 | FBgn0035496 | Serine protease homolog 35 |
| LG2 | 6361730 | TC030253 | 0 | intron | Gustatory receptor | - | - |
| LG2 | 6361884 | TC000829 | 0 | exon | Serine protease H18 | FBgn0035496 | Serine protease homolog 35 |
| LG2 | 6361884 | TC030253 | 0 | intron | Gustatory receptor | - | - |
| LG2 | 6362638 | TC000829 | 0 | intron | Serine protease H18 | FBgn0035496 | Serine protease homolog 35 |
| LG2 | 6363263 | TC000829 | 0 | intron | " | " | " |
| LG2 | 6363541 | TC000829 | 0 | intron | " | " | " |
| LG2 | 6363803 | TC000829 | 0 | intron | " | " | " |
| LG2 | 6363874 | TC000829 | 0 | intron | " | " | " |
| LG2 | 6364637 | TC000829 | 0 | intron | " | " | " |
| LG2 | 6364854 | TC000829 | 0 | intron | " | " | " |
| LG2 | 6364859 | TC000829 | 0 | intron | " | " | " |
| LG2 | 6366349 | TC000829 | 298 | - | " | " | " |
| LG2 | 6366418 | TC000829 | 367 | - | " | " | " |
| LG2 | 6367320 | TC000829 | 1269 | - | " | " | " |
| LG2 | 6367492 | TC000829 | 1441 | - | " | " | " |
| LG2 | 6367750 | TC000829 | 1699 | - | " | " | " |

| | | | | | | | |
|---|---|---|---|---|---|---|---|
| LG2 | 6367753 | TC000829 | 1702 | - | " | " | " |
| LG2 | 6367763 | TC000829 | 1712 | - | " | " | " |
| LG2 | 6367974 | TC000829 | 1923 | - | " | " | " |
| LG2 | 6368123 | TC000829 | 2072 | - | " | " | " |
| LG2 | 6368173 | TC000829 | 2122 | - | " | " | " |
| LG2 | 6369222 | TC000829 | 3171 | - | " | " | " |
| LG2 | 6369242 | TC000829 | 3191 | - | " | " | " |
| LG2 | 6369280 | TC000829 | 3229 | - | " | " | " |
| LG2 | 6371431 | TC000829 | 5380 | - | " | " | " |
| LG2 | 6371729 | TC000829 | 5678 | - | " | " | " |
| LG2 | 6371733 | TC000829 | 5682 | - | " | " | " |
| LG2 | 6371741 | TC000829 | 5690 | - | " | " | " |
| LG2 | 6371746 | TC000829 | 5695 | - | " | " | " |
| LG2 | 6371749 | TC000829 | 5698 | - | " | " | " |
| LG2 | 6372049 | TC000829 | 5998 | - | " | " | " |
| LG2 | 6372237 | TC000829 | 6186 | - | " | " | " |
| LG2 | 6372297 | TC000829 | 6246 | - | " | " | " |
| LG2 | 6373869 | TC000829 | 7818 | - | " | " | " |
| LG2 | 6759639 | TC032076 | 0 | intron | Disks large 1 tumor suppressor protein-like Protein | FBgn0001624 | discs large 1 |
| LG2 | 6759675 | TC032076 | 0 | intron | " | " | " |
| LG2 | 6767618 | TC032076 | 0 | intron | " | " | " |
| LG2 | 12685100 | TC001207 | 0 | intron | RNA-binding protein MEX3B-like Protein | FBgn0039920 | CG11360 |
| LG2 | 12695008 | TC032210 | 1005 | - | - | - | - |
| LG2 | 12695027 | TC032210 | 1024 | - | - | - | - |
| LG2 | 12695032 | TC032210 | 1029 | - | - | - | - |
| LG2 | 12696786 | TC000351 | 1162 | - | - | - | - |
| LG2 | 12708148 | TC000351 | 0 | intron | - | - | - |

| | | | | | | | |
|---|---|---|---|---|---|---|---|
| LG2 | 12710288 | TC000351 | 0 | intron | - | - | - |
| LG2 | 12710438 | TC000351 | 0 | intron | - | - | - |
| LG2 | 12712223 | TC000351 | 0 | intron | - | - | - |
| LG3 | 25378890 | TC034629 | 0 | intron | Hepatic leukemia factor-like protein | FBgn0016694 | PAR-domain protein 1 |
| LG3 | 25383208 | TC034629 | 0 | intron | " | " | " |
| LG3 | 25385605 | TC034629 | 0 | intron | " | " | " |
| LG3 | 25386766 | TC034629 | 0 | intron | " | " | " |
| LG4 | 9239595 | TC008372 | 0 | intron | Leucine--tRNA ligase%2C cytoplasmic-like Protein | FBgn0284253 | Leucyl-tRNA synthetase |
| LG4 | 9239957 | TC008372 | 0 | intron | " | " | " |
| LG4 | 9240001 | TC008372 | 0 | intron | " | " | " |
| LG4 | 9240050 | TC008372 | 0 | intron | " | " | " |
| LG4 | 9240163 | TC008372 | 0 | intron | " | " | " |
| LG4 | 9240232 | TC008372 | 0 | intron | " | " | " |
| LG4 | 9240233 | TC008372 | 0 | intron | " | " | " |
| LG4 | 9240250 | TC008372 | 0 | intron | " | " | " |
| LG4 | 9240287 | TC008372 | 0 | intron | " | " | " |
| LG4 | 9240308 | TC008372 | 0 | intron | " | " | " |
| LG4 | 9240313 | TC008372 | 0 | intron | " | " | " |
| LG4 | 9240325 | TC008372 | 0 | intron | " | " | " |
| LG4 | 9240394 | TC008372 | 0 | intron | " | " | " |
| LG4 | 9248921 | TC007110 | 1542 | - | - | FBgn0031016 | kekkon 5 |
| LG4 | 9248923 | TC007110 | 1540 | - | " | " | " |
| LG4 | 9248925 | TC007110 | 1538 | - | " | " | " |
| LG6 | 9546142 | TC014863 | 2200 | - | 43 kDa receptor-associated protein of the synapse homolog-like Protein | FBgn0039911 | CG1909 |
| LG6 | 9620940 | TC014861 | 0 | exon | Neuroblastoma%2C suppression of tumorigenicity 1 | - | - |
| LG6 | 9621142 | TC014861 | 0 | exon | " | - | - |

| | | | | | | | |
|---|---|---|---|---|---|---|---|
| LG6 | 9623478 | TC014861 | 0 | intron | " | - | - |
| LG6 | 9623731 | TC014861 | 0 | exon | " | - | - |
| LG6 | 9625178 | TC014861 | 0 | intron | " | - | - |
| LG6 | 9632402 | TC014860 | 0 | intron | Moesin/ezrin/radixin homolog 2-like Protein | FBgn0086384 | Merlin |
| LG6 | 9632454 | TC014860 | 0 | intron | " | " | " |
| LG6 | 9640445 | TC014859 | 0 | intron | Zinc finger HIT domain-containing protein 2-like Protein | FBgn0051223 | CG31223 |
| LG6 | 9640628 | TC014859 | 0 | intron | " | " | " |
| LG6 | 9641505 | TC014859 | 0 | intron | " | " | " |
| LG6 | 9641518 | TC014859 | 0 | intron | " | " | " |
| LG6 | 9647714 | TC014857 | 0 | exon | Ribosome biogenesis methyltransferase WBSCR22-like Protein | FBgn0037543 | CG10903 |
| LG6 | 9648497 | TC015858 | 37 | - | Cell division control protein 45 homolog-like Protein | FBgn0026143 | Cell division cycle 45 |
| LG6 | 9648756 | TC015858 | 0 | exon | " | " | " |
| LG6 | 9649654 | TC015858 | 0 | exon | " | " | " |
| LG6 | 9649716 | TC015858 | 0 | intron | " | " | " |
| LG6 | 9649822 | TC015858 | 0 | intron | " | " | " |
| LG6 | 9649865 | TC015858 | 0 | intron | " | " | " |
| LG6 | 9652242 | TC015858 | 0 | exon | " | " | " |
| LG6 | 9652269 | TC015858 | 0 | exon | " | " | " |
| LG6 | 9652392 | TC015858 | 0 | exon | " | " | " |
| LG6 | 9659272 | TC014856 | 0 | intron | - | - | - |
| LG6 | 9662151 | TC014856 | 0 | exon | - | - | - |
| LG6 | 9664520 | TC014856 | 0 | intron | - | - | - |
| LG6 | 9664534 | TC014856 | 0 | intron | - | - | - |
| LG6 | 9664921 | TC014856 | 0 | intron | - | - | - |
| LG6 | 9665270 | TC014856 | 0 | intron | - | - | - |
| LG6 | 9669740 | TC014856 | 0 | intron | - | - | - |

| | | | | | | | |
|---|---|---|---|---|---|---|---|
| LG6 | 9669776 | TC014856 | 0 | intron | - | - | - |
| LG6 | 9670076 | TC014856 | 0 | intron | - | - | - |
| LG6 | 9671781 | TC014856 | 0 | intron | - | - | - |
| LG6 | 9674940 | TC014856 | 0 | intron | - | - | - |
| LG6 | 9675271 | TC014856 | 0 | intron | - | - | - |
| LG6 | 9675299 | TC014856 | 0 | intron | - | - | - |
| LG6 | 9675442 | TC014856 | 0 | intron | - | - | - |
| LG6 | 9686696 | TC014856 | 5404 | - | - | - | - |
| LG6 | 9686813 | TC014856 | 5521 | - | - | - | - |
| LG7 | 1184860 | TC009377 | 776 | - | - | FBgn0083951 | CG34115 |
| LG7 | 1184872 | TC009377 | 764 | - | - | " | " |
| LG7 | 1184873 | TC009377 | 763 | - | - | " | " |
| LG7 | 1188869 | TC009441 | 0 | intron | Collagen type IV alpha-3-binding protein-like Protein | FBgn0027569 | ceramide transfer protein |
| LG8 | 12415127 | TC005479 | 0 | intron | Polypeptide N-acetylgalactosaminyltransferase | FBgn0050463 | Polypeptide N-Acetylgalactosaminyltransferase 9 |
| LG8 | 12415169 | TC005479 | 0 | intron | " | " | " |
| LG8 | 12415200 | TC005479 | 0 | intron | " | " | " |
| LG8 | 12415247 | TC005479 | 0 | intron | " | " | " |
| LG8 | 12415408 | TC005479 | 0 | intron | " | " | " |
| LG8 | 12459002 | TC006646 | 2969 | - | - | FBgn0038819 | Cuticular protein 92F |
| LG8 | 12459035 | TC006646 | 2936 | - | - | " | " |
| LG8 | 12509462 | TC005477 | 3 | - | Serine/threonine-protein kinase ULK2-like Protein | FBgn0260945 | Autophagy-related 1 |
| LG8 | 12509497 | TC005477 | 38 | - | " | " | " |
| LG8 | 12511174 | TC034152 | 0 | 3` UTR | - | - | - |
| LG8 | 12538848 | TC034154 | 508 | - | - | - | - |
| LG8 | 12539021 | TC034154 | 681 | - | - | - | - |
| LG8 | 12540431 | TC034154 | 2091 | - | - | - | - |

| | | | | | | | |
|---|---|---|---|---|---|---|---|
| LG8 | 12545606 | TC034154 | 7266 | - | - | - | - |
| LG8 | 12545612 | TC034154 | 7272 | - | - | - | - |
| LG8 | 12545951 | TC034154 | 7611 | - | - | - | - |
| LG8 | 12565644 | TC005473 | 0 | intron | - | FBgn0263994 | CG43737 |
| LG8 | 12569527 | TC005473 | 0 | intron | - | " | " |
| LG9 | 8097869 | TC034351 | 0 | exon | - | FBgn0261836 | Muscle-specific protein 300 kDa |
| LG9 | 8103170 | TC034351 | 0 | exon | - | " | " |
| LG9 | 8103173 | TC034351 | 0 | exon | - | " | " |
| LG9 | 8113123 | TC034351 | 0 | exon | - | " | " |
| LG9 | 8804148 | TC034357 | 18651 | - | - | FBgn0287478 | Calmodulin-binding transcription activator |
| LG9 | 8805772 | TC034357 | 17027 | - | - | " | " |
| LG9 | 8815857 | TC034357 | 6942 | - | - | " | " |
| LG9 | 8815958 | TC034357 | 6841 | - | - | " | " |
| LG9 | 8817105 | TC034357 | 5694 | - | - | " | " |
| LG9 | 8817652 | TC034357 | 5147 | - | - | " | " |
| LG9 | 8822755 | TC034357 | 44 | - | - | " | " |
| LG9 | 8824374 | TC034357 | 0 | 3`UTR | - | " | " |
| LG9 | 8824863 | TC034357 | 0 | intron | - | " | " |
| LG9 | 8825322 | TC034357 | 0 | intron | - | " | " |
| LG9 | 8830110 | TC034357 | 0 | intron | - | " | " |
| LG9 | 8830151 | TC034357 | 0 | intron | - | " | " |
| LG9 | 8831762 | TC034357 | 0 | intron | - | " | " |
| LG9 | 8834324 | TC034357 | 0 | intron | - | " | " |
| LG9 | 8838455 | TC034357 | 0 | intron | - | " | " |
| LG9 | 8839515 | TC034357 | 0 | intron | - | " | " |
| LG9 | 8839675 | TC034357 | 0 | intron | - | " | " |
| LG9 | 8839823 | TC034357 | 0 | intron | - | " | " |

| | | | | | | | |
|---|---|---|---|---|---|---|---|
| LG9 | 8839890 | TC034357 | 0 | intron | - | " | " |
| LG9 | 8839917 | TC034357 | 0 | intron | - | " | " |
| LG9 | 8839929 | TC034357 | 0 | intron | - | " | " |
| LG9 | 8839973 | TC034357 | 0 | intron | - | " | " |
| LG9 | 8839977 | TC034357 | 0 | intron | - | " | " |
| LG9 | 8839991 | TC034357 | 0 | intron | - | " | " |
| LG9 | 8839994 | TC034357 | 0 | intron | - | " | " |
| LG9 | 8840023 | TC034357 | 0 | intron | - | " | " |
| LG9 | 8840043 | TC034357 | 0 | intron | - | " | " |
| LG9 | 8840087 | TC034357 | 0 | intron | - | " | " |
| LG9 | 8840169 | TC034357 | 0 | intron | - | " | " |
| LG9 | 8840559 | TC034357 | 0 | exon | - | " | " |
| LG9 | 8840560 | TC034357 | 0 | exon | - | " | " |
| LG9 | 8840575 | TC034357 | 0 | exon | - | " | " |
| LG9 | 8843888 | TC034357 | 0 | intron | - | " | " |
| LG9 | 8844155 | TC034357 | 0 | intron | - | " | " |
| LG9 | 8844190 | TC034357 | 0 | intron | - | " | " |
| LG9 | 8844302 | TC034357 | 0 | intron | - | " | " |
| LG9 | 8844346 | TC034357 | 0 | intron | - | " | " |
| LG9 | 8844347 | TC034357 | 0 | intron | - | " | " |
| LG9 | 8844348 | TC034357 | 0 | intron | - | " | " |
| LG9 | 8844460 | TC034357 | 0 | intron | - | " | " |
| LG9 | 8844465 | TC034357 | 0 | intron | - | " | " |
| LG9 | 8844499 | TC034357 | 0 | intron | - | " | " |
| LG9 | 8844514 | TC034357 | 0 | intron | - | " | " |
| LG9 | 8844602 | TC034357 | 0 | intron | - | " | " |
| LG9 | 8844715 | TC034357 | 0 | intron | - | " | " |
| LG9 | 8848978 | TC034357 | 0 | intron | - | " | " |

| | | | | | | | |
|---|---|---|---|---|---|---|---|
| LG9 | 8849778 | TC034357 | 0 | intron | - | " | " |
| LG9 | 8850259 | TC034357 | 0 | intron | - | " | " |
| LG9 | 8850303 | TC034357 | 0 | intron | - | " | " |
| LG9 | 8851956 | TC034357 | 0 | intron | - | " | " |
| LG9 | 8853079 | TC034357 | 0 | intron | - | " | " |
| LG9 | 8853087 | TC034357 | 0 | intron | - | " | " |
| LG9 | 8853094 | TC034357 | 0 | intron | - | " | " |
| LG9 | 8871126 | TC034357 | 0 | intron | - | " | " |
| LG9 | 8871142 | TC034357 | 0 | intron | - | " | " |
| LG9 | 8872540 | TC034357 | 0 | intron | - | " | " |
| LG9 | 8872611 | TC034357 | 0 | intron | - | " | " |
| LG9 | 8872613 | TC034357 | 0 | intron | - | " | " |
| LG9 | 8872659 | TC034357 | 0 | intron | - | " | " |
| LG9 | 8881740 | TC034357 | 0 | intron | - | " | " |
| LGX | 3211320 | TC014210 | 28 | - | Guanylate kinase-like Protein | FBgn0036099 | oya |
| LGX | 3211495 | TC014210 | 0 | 5' UTR | " | " | " |
| LGX | 3211514 | TC014210 | 0 | 5' UTR | " | " | " |
| LGX | 3211671 | TC014210 | 0 | intron | " | " | " |
| LGX | 3212007 | TC014210 | 0 | exon | " | " | " |
| LGX | 3212208 | TC014210 | 0 | intron | " | " | " |
| LGX | 3212306 | TC014210 | 0 | exon | " | " | " |
| LGX | 3212954 | TC014210 | 330 | - | " | " | " |
| LGX | 3212987 | TC014210 | 363 | - | " | " | " |
| LGX | 3213547 | TC013603 | 330 | - | Gamma-glutamylcyclotransferase | FBgn0030411 | CG2540 |
| LGX | 3213563 | TC013603 | 314 | - | " | " | " |
| LGX | 3231380 | TC013602 | 3409 | - | Vitellogenin-like Protein | - | - |
| LGX | 3231485 | TC013602 | 3304 | - | " | - | - |
| LGX | 3234141 | TC013602 | 648 | - | " | - | - |

| | | | | | | | |
|---|---|---|---|---|---|---|---|
| LGX | 3234628 | TC013602 | 161 | - | " | - | - |
| LGX | 3234672 | TC013602 | 117 | - | " | - | - |
| LGX | 3234990 | TC013602 | 0 | exon | " | - | - |
| LGX | 3235611 | TC013602 | 0 | exon | " | - | - |
| LGX | 3236513 | TC013602 | 0 | exon | " | - | - |
| LGX | 3236534 | TC013602 | 0 | exon | " | - | - |
| LGX | 3236618 | TC013602 | 0 | exon | " | - | - |
| LGX | 3236786 | TC013602 | 0 | exon | " | - | - |
| LGX | 3236915 | TC013602 | 0 | exon | " | - | - |
| LGX | 3237068 | TC013602 | 0 | exon | " | - | - |
| LGX | 3237077 | TC013602 | 0 | exon | " | - | - |
| LGX | 3237179 | TC013602 | 0 | exon | " | - | - |
| LGX | 3237272 | TC013602 | 0 | exon | " | - | - |
| LGX | 3237278 | TC013602 | 0 | exon | " | - | - |
| LGX | 3237305 | TC013602 | 0 | exon | " | - | - |
| LGX | 3237479 | TC013602 | 0 | exon | " | - | - |
| LGX | 3237795 | TC013602 | 0 | exon | " | - | - |
| LGX | 3238015 | TC013602 | 0 | exon | " | - | - |
| LGX | 3238276 | TC013602 | 0 | exon | " | - | - |
| LGX | 3238312 | TC013602 | 0 | exon | " | - | - |
| LGX | 3238330 | TC013602 | 0 | exon | " | - | - |
| LGX | 3238365 | TC013602 | 0 | exon | " | - | - |
| LGX | 3239065 | TC013602 | 0 | exon | " | - | - |
| LGX | 3239137 | TC013602 | 0 | exon | " | - | - |
| LGX | 3239356 | TC013602 | 0 | exon | " | - | - |
| LGX | 3239398 | TC013602 | 0 | exon | " | - | - |
| LGX | 3240529 | TC013602 | 43 | - | " | - | - |
| LGX | 3241290 | TC014211 | 0 | 5' UTR | Neuropeptide Y receptor-like Protein | FBgn0038880 | SIFamide receptor |

| | | | | | | | |
|---|---|---|---|---|---|---|---|
| LGX | 3242080 | TC014211 | 0 | intron | " | " | " |
| LGX | 3242093 | TC014211 | 0 | intron | " | " | " |
| LGX | 3242120 | TC014211 | 0 | intron | " | " | " |
| LGX | 3242820 | TC014211 | 0 | intron | " | " | " |
| LGX | 3242897 | TC014211 | 0 | exon | " | " | " |
| LGX | 3243399 | TC014211 | 0 | intron | " | " | " |
| LGX | 3243456 | TC014211 | 0 | intron | " | " | " |
| LGX | 3243543 | TC014211 | 0 | intron | " | " | " |
| LGX | 3244121 | TC014211 | 0 | exon | " | " | " |
| LGX | 3244294 | TC014211 | 0 | exon | " | " | " |
| LGX | 3244641 | TC014211 | 81 | - | " | " | " |
| LGX | 3244850 | TC014211 | 290 | - | " | " | " |
| LGX | 3245312 | TC013601 | 367 | - | Putative DNA mismatch repair protein Msh6-like Protein | FBgn0036486 | Msh6 |
| LGX | 3248338 | TC013601 | 0 | intron | " | " | " |
| LGX | 3248353 | TC013601 | 0 | intron | " | " | " |
| LGX | 3248379 | TC013601 | 0 | intron | " | " | " |
| LGX | 3248670 | TC013601 | 0 | intron | " | " | " |
| LGX | 3248686 | TC013601 | 0 | intron | " | " | " |
| LGX | 3248697 | TC013601 | 0 | intron | " | " | " |
| LGX | 3248783 | TC013601 | 0 | intron | " | " | " |
| LGX | 3248929 | TC013601 | 0 | intron | " | " | " |
| LGX | 3256119 | TC031503 | 0 | intron | Xaa-Pro dipeptidase-like Protein | FBgn0000455 | Dipeptidase C |
| LGX | 3257072 | TC031503 | 0 | intron | " | " | " |
| LGX | 3257317 | TC031503 | 0 | intron | " | " | " |
| LGX | 3257872 | TC031503 | 0 | intron | " | " | " |
| LGX | 3257880 | TC031503 | 0 | intron | " | " | " |
| LGX | 3258590 | TC031503 | 0 | intron | " | " | " |

| | | | | | | | |
|---|---|---|---|---|---|---|---|
| LGX | 3258917 | TC031503 | 0 | intron | " | " | " |
| LGX | 3259209 | TC031503 | 0 | intron | " | " | " |
| LGX | 3259736 | TC031503 | 0 | intron | " | " | " |
| LGX | 3259806 | TC031503 | 0 | intron | " | " | " |
| LGX | 3262032 | TC031503 | 0 | intron | " | " | " |
| LGX | 3262904 | TC031503 | 0 | intron | " | " | " |
| LGX | 3285248 | TC030882 | 0 | intron | Arginine kinase-like Protein | FBgn0000116 | Arginine kinase 1 |
| LGX | 3291587 | TC031505 | 0 | intron | - | - | - |
| LGX | 3291733 | TC031505 | 0 | intron | - | - | - |
| LGX | 3297041 | TC013595 | 0 | 5' UTR | - | FBgn0030847 | CG12991 |
| LGX | 3410074 | TC013586 | 0 | intron | - | FBgn0261885 | osa |
| LGX | 3410669 | TC013586 | 0 | intron | - | " | " |
| LGX | 3410880 | TC013586 | 0 | intron | - | " | " |
| LGX | 3410895 | TC013586 | 0 | intron | - | " | " |
| LGX | 3411038 | TC013586 | 0 | intron | - | " | " |
| LGX | 3411150 | TC013586 | 0 | intron | - | " | " |
| LGX | 3412330 | TC013586 | 0 | intron | - | " | " |
| LGX | 3413050 | TC013586 | 0 | intron | - | " | " |
| LGX | 3413380 | TC013586 | 0 | intron | - | " | " |
| LGX | 3413719 | TC013586 | 0 | intron | - | " | " |
| LGX | 3413791 | TC013586 | 0 | intron | - | " | " |
| LGX | 3413897 | TC013586 | 0 | intron | - | " | " |
| LGX | 3414129 | TC013586 | 0 | intron | - | " | " |
| LGX | 3414474 | TC013586 | 0 | intron | - | " | " |
| LGX | 3416432 | TC013586 | 0 | intron | - | " | " |
| LGX | 3416597 | TC013586 | 0 | intron | - | " | " |
| LGX | 3416687 | TC013586 | 0 | intron | - | " | " |
| LGX | 3417628 | TC013586 | 0 | intron | - | " | " |

| | | | | | | | |
|---|---|---|---|---|---|---|---|
| LGX | 3417908 | TC013586 | 0 | intron | - | " | " |
| LGX | 3417923 | TC013586 | 0 | intron | - | " | " |
| LGX | 3418078 | TC013586 | 0 | intron | - | " | " |
| LGX | 3418168 | TC013586 | 0 | intron | - | " | " |
| LGX | 3418294 | TC013586 | 0 | intron | - | " | " |
| LGX | 3418757 | TC013586 | 0 | intron | - | " | " |
| LGX | 3418817 | TC013586 | 0 | intron | - | " | " |
| LGX | 3418900 | TC013586 | 0 | intron | - | " | " |
| LGX | 3419421 | TC013586 | 0 | intron | - | " | " |
| LGX | 3419477 | TC013586 | 0 | intron | - | " | " |
| LGX | 3419563 | TC013586 | 0 | intron | - | " | " |
| LGX | 3421349 | TC013586 | 0 | intron | - | " | " |
| LGX | 3421484 | TC013586 | 0 | intron | - | " | " |
| LGX | 3421675 | TC013586 | 0 | intron | - | " | " |
| LGX | 3422074 | TC013586 | 0 | intron | - | " | " |
| LGX | 3422103 | TC013586 | 0 | intron | - | " | " |
| LGX | 3422156 | TC013586 | 0 | intron | - | " | " |
| LGX | 3422270 | TC013586 | 0 | intron | - | " | " |
| LGX | 3424571 | TC013586 | 0 | 5' UTR | - | " | " |
| LGX | 3425229 | TC013585 | 0 | 3' UTR | Katanin p60 ATPase-containing subunit A1 | FBgn0040208 | Katanin 60 |
| LGX | 3425246 | TC013585 | 0 | 3' UTR | " | " | " |
| LGX | 3425252 | TC013585 | 0 | 3' UTR | " | " | " |
| LGX | 3425366 | TC013585 | 0 | 3' UTR | " | " | " |
| LGX | 3425615 | TC013585 | 0 | exon | " | " | " |
| LGX | 3426178 | TC013585 | 0 | exon | " | " | " |
| LGX | 3426416 | TC013585 | 0 | exon | " | " | " |
| LGX | 3426671 | TC013585 | 0 | intron | " | " | " |
| LGX | 3426782 | TC013585 | 0 | exon | " | " | " |

| | | | | | | | |
|---|---|---|---|---|---|---|---|
| LGX | 3426791 | TC013585 | 0 | exon | " | " | " |
| LGX | 3426927 | TC013585 | 0 | intron | " | " | " |
| LGX | 3426989 | TC013585 | 0 | exon | " | " | " |
| LGX | 3427004 | TC013585 | 0 | exon | " | " | " |
| LGX | 3427037 | TC013585 | 0 | exon | " | " | " |
| LGX | 3427266 | TC013585 | 0 | - | " | " | " |
| LGX | 3427580 | TC013585 | 28 | - | " | " | " |
| LGX | 3427596 | TC013585 | 44 | - | " | " | " |
| LGX | 3427610 | TC013585 | 58 | - | " | " | " |
| LGX | 3427632 | TC013585 | 80 | - | " | " | " |
| LGX | 3427664 | TC013585 | 112 | - | " | " | " |
| LGX | 3427704 | TC014218 | 120 | - | RNA pseudouridylate synthase domain-containing protein 1-like Protein | - | - |
| LGX | 3427739 | TC014218 | 85 | - | " | - | - |
| LGX | 3427745 | TC014218 | 79 | - | " | - | - |
| LGX | 3427895 | TC014218 | 0 | exon | " | - | - |
| LGX | 3427908 | TC014218 | 0 | exon | " | - | - |
| LGX | 3428246 | TC014218 | 0 | exon | " | - | - |
| LGX | 3428338 | TC014218 | 0 | intron | " | - | - |
| LGX | 3428391 | TC014218 | 0 | exon | " | - | - |
| LGX | 3428639 | TC014218 | 0 | intron | " | - | - |
| LGX | 3428822 | TC014218 | 0 | 3' UTR | " | - | - |
| LGX | 3429228 | TC014219 | 0 | exon | Protein jagunal-like Protein | FBgn0037374 | jagunal |
| LGX | 3429573 | TC014219 | 0 | intron | " | " | " |
| LGX | 3429575 | TC014219 | 0 | intron | " | " | " |
| LGX | 3429711 | TC014219 | 0 | exon | " | " | " |
| LGX | 3429733 | TC014219 | 0 | 3' UTR | " | " | " |
| LGX | 3429815 | TC014219 | 0 | 3' UTR | " | " | " |

| | | | | | | | |
|---|---|---|---|---|---|---|---|
| LGX | 3430593 | TC014220 | 0 | intron | Monocarboxylate transporter 9-like Protein | FBgn0033028 | hermes |
| LGX | 3430846 | TC014220 | 0 | intron | " | " | " |
| LGX | 3430854 | TC014220 | 0 | intron | " | " | " |
| LGX | 3431112 | TC014220 | 0 | 5' UTR | " | " | " |
| LGX | 3431113 | TC014220 | 0 | 5' UTR | " | " | " |
| LGX | 3431117 | TC014220 | 0 | 5' UTR | " | " | " |
| LGX | 3432141 | TC014220 | 0 | intron | " | " | " |
| LGX | 3432185 | TC014220 | 0 | intron | " | " | " |
| LGX | 3432202 | TC014220 | 0 | intron | " | " | " |
| LGX | 3432226 | TC014220 | 0 | intron | " | " | " |
| LGX | 3432236 | TC014220 | 0 | intron | " | " | " |
| LGX | 3432259 | TC014220 | 0 | intron | " | " | " |
| LGX | 3432728 | TC014220 | 0 | intron | " | " | " |
| LGX | 3432769 | TC014220 | 0 | intron | " | " | " |
| LGX | 3432931 | TC014220 | 0 | exon | " | " | " |
| LGX | 3433392 | TC014220 | 0 | exon | " | " | " |
| LGX | 3433495 | TC014220 | 0 | exon | " | " | " |
| LGX | 3434350 | TC014220 | 0 | 3' UTR | " | " | " |
| LGX | 3434839 | TC014220 | 116 | - | " | " | " |
| LGX | 3435444 | TC014221 | 0 | intron | UPF0547 protein C16orf87 homolog-like Protein | - | - |
| LGX | 3435653 | TC014221 | 0 | exon | " | - | - |
| LGX | 3435837 | TC014221 | 0 | 3' UTR | " | - | - |
| LGX | 3435837 | TC013584 | 0 | intron | - | - | - |
| LGX | 3435838 | TC014221 | 0 | 3' UTR | UPF0547 protein C16orf87 homolog-like Protein | - | - |
| LGX | 3435838 | TC013584 | 0 | intron | - | - | - |
| LGX | 3435883 | TC014221 | 0 | 3' UTR | UPF0547 protein C16orf87 homolog-like Protein | - | - |

| | | | | | | | |
|---|---|---|---|---|---|---|---|
| LGX | 3435883 | TC013584 | 0 | intron | - | - | - |
| LGX | 3435884 | TC014221 | 0 | 3' UTR | UPF0547 protein C16orf87 homolog-like Protein | - | - |
| LGX | 3435884 | TC013584 | 0 | intron | - | - | - |
| LGX | 3435891 | TC014221 | 0 | 3' UTR | UPF0547 protein C16orf87 homolog-like Protein | - | - |
| LGX | 3435891 | TC013584 | 0 | intron | - | - | - |
| LGX | 3436008 | TC014221 | 0 | 3' UTR | UPF0547 protein C16orf87 homolog-like Protein | - | - |
| LGX | 3436008 | TC013584 | 0 | intron | - | - | - |
| LGX | 3436353 | TC014222 | 0 | 5' UTR | Nischarin-like Protein | FBgn0033996 | CG11807 |
| LGX | 3436864 | TC014222 | 0 | intron | " | " | " |
| LGX | 3437510 | TC014222 | 0 | exon | " | " | " |
| LGX | 3437580 | TC014222 | 0 | exon | " | " | " |
| LGX | 3437916 | TC014222 | 0 | exon | " | " | " |
| LGX | 3438014 | TC014222 | 0 | 3' UTR | " | " | " |
| LGX | 3438066 | TC014222 | 49 | - | " | " | " |
| LGX | 3438498 | TC030883 | 0 | 5' UTR | - | - | - |
| LGX | 3438602 | TC030883 | 0 | intron | - | - | - |
| LGX | 3438812 | TC030883 | 0 | 5' UTR | - | - | - |
| LGX | 3438871 | TC030883 | 0 | exon | - | - | - |
| LGX | 3438873 | TC030883 | 0 | exon | - | - | - |
| LGX | 3439103 | TC030883 | 0 | intron | - | - | - |
| LGX | 3439637 | TC030883 | 0 | intron | - | - | - |
| LGX | 3439666 | TC030883 | 0 | intron | - | - | - |
| LGX | 3439708 | TC030883 | 0 | intron | - | - | - |
| LGX | 3439777 | TC030883 | 0 | intron | - | - | - |
| LGX | 3439851 | TC030883 | 0 | intron | - | - | - |
| LGX | 3439955 | TC030883 | 0 | intron | - | - | - |

| | | | | | | | |
|---|---|---|---|---|---|---|---|
| LGX | 3440721 | TC030883 | 0 | exon | - | - | - |
| LGX | 3440934 | TC030883 | 67 | - | - | - | - |
| LGX | 3441341 | TC030884 | 32 | - | Phosphatidylinositol 3-kinase regulatory subunit alpha-like Protein | FBgn0020622 | Pi3K21B |
| LGX | 3444819 | TC030884 | 0 | intron | " | " | " |
| LGX | 3444894 | TC030884 | 0 | intron | " | " | " |
| LGX | 3445814 | TC030884 | 0 | intron | " | " | " |
| LGX | 3446359 | TC030884 | 0 | intron | " | " | " |
| LGX | 3449862 | TC030884 | 0 | intron | " | " | " |
| LGX | 3449981 | TC030884 | 0 | intron | " | " | " |
| LGX | 3453755 | TC030884 | 0 | exon | " | " | " |
| LGX | 3456897 | TC013582 | 0 | exon | Cathepsin K | FBgn0032228 | Cathepsin L4 |
| LGX | 3457479 | TC013582 | 0 | exon | " | | |
| LGX | 3457721 | TC014225 | 0 | 5'UTR | Transcription factor E3-like Protein | FBgn0263112 | Mitf |
| LGX | 3457804 | TC014225 | 0 | 5' UTR | " | " | " |
| LGX | 3460463 | TC014225 | 0 | intron | " | " | " |
| LGX | 3464581 | TC014225 | 0 | intron | " | " | " |
| LGX | 3465929 | TC014225 | 0 | intron | " | " | " |
| LGX | 3466970 | TC014225 | 0 | intron | " | " | " |
| LGX | 3467613 | TC014225 | 0 | intron | " | " | " |
| LGX | 3468887 | TC014225 | 0 | 3' UTR | " | " | " |
| LGX | 3468975 | TC014225 | 0 | 3' UTR | " | " | " |
| LGX | 3470039 | TC014225 | 925 | - | " | " | " |
| LGX | 3470189 | TC014225 | 1075 | - | " | " | " |
| LGX | 3470239 | TC014225 | 1125 | - | " | " | " |
| LGX | 3470249 | TC014225 | 1135 | - | " | " | " |
| LGX | 3471429 | TC014226 | 321 | - | Kinesin-like protein KIF13A | FBgn0019968 | Kinesin heavy chain 73 |
| LGX | 3472717 | TC014226 | 0 | intron | " | " | " |

| | | | | | | | |
|---|---|---|---|---|---|---|---|
| LGX | 3473260 | TC014226 | 0 | intron | " | " | " |

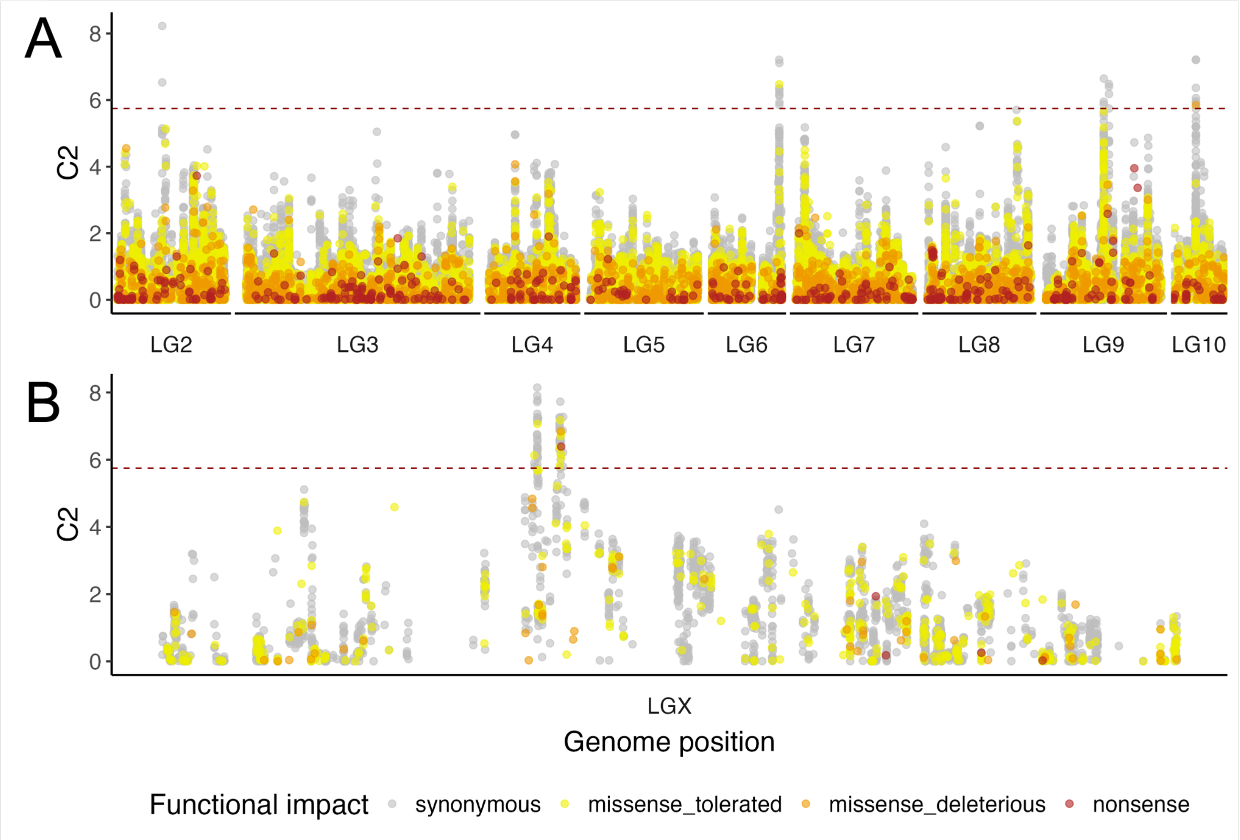

**Figure S7.** Genomic loci showing allele frequency differentiation associated with sexual selection regime in experimentally evolving Tribolium castaneum populations, coloured by functional impact. Values of the C2 statistic generated by BayPass analysis (Olazcuaga et al. 2020) measuring SNP-wise associations with the two sexual selection regimes (polyandrous vs. monandrous) while accounting for shared population history. The dotted line shows the 0.999 quantile of C2 values from a BayPass run on a pseudo-observed dataset of putatively neutral SNPs. Colours denote the functional impact derived via protein logic (SnpEff; Cingolani et al. 2012) and conservance score for non-synonymous variants (SIFT; Vaser et al. 2016). Separate analyses were performed for **A)** autosomal linkage groups, and **B)** the X chromosome (LGX).
